# Dynamic expression and localization of the LIN-2/7/10 protein scaffolding complex during *C. elegans* vulval development

**DOI:** 10.1101/2020.06.17.157958

**Authors:** Kimberley D. Gauthier, Christian E. Rocheleau

**Affiliations:** Division of Endocrinology and Metabolism, Department of Medicine, and Department of Anatomy and Cell Biology, McGill University; and the Metabolic Disorders and Complications Program, Centre for Translational Biology, Research Institute of the McGill University Health Centre, Montreal, QC H4A 3J1, Canada

**Author notes:** Author to whom correspondence should be addressed, Christian Rocheleau, Research Institute of the McGill University Health Centre, 1001 Décarie Blvd, Room E02.7242, Montreal, QC, H4A 3J1.

**Keywords:** *Caenorhabditis elegans*, vulva, LET-23, Cask, Lin7, APBA

## Abstract

The evolutionarily conserved LIN-2 (CASK)/LIN-7 (Lin7A-C)/LIN-10 (APBA1) complex plays an important role in regulating spatial organization of membrane proteins and signaling components. In *C. elegans,* the complex is essential for development of the vulva by promoting the localization of the sole Epidermal Growth Factor Receptor (EGFR) orthologue, LET-23, to the basolateral membrane of the vulva precursor cells (VPCs) where it can specify the vulval cell fate. However, the expression and localization of the LIN-2/7/10 complex, and how the complex regulates receptor localization, are not known. Here we describe an *in vivo* analysis of the complex in *C. elegans* VPCs. Only LIN-7 colocalizes with LET-23 EGFR at the basolateral membrane, while the LIN-2/7/10 complex components instead colocalize at cytoplasmic foci, consistent with Golgi or endosomes. LIN-10 recruits LIN-2, which in turn recruits LIN-7. We demonstrate that the complex forms *in vivo* with particularly strong interaction and colocalization between LIN-2 and LIN-7. Our data suggest that the LIN-2/7/10 complex forms on endomembrane compartments where it likely targets LET-23 EGFR to the basolateral membrane, and point to distinct regulation between LIN-2/7 and LIN-10.

**Summary Statement:** LIN-10 recruits LIN-2 and LIN-7 to Golgi or recycling endosomes, consistent with targeting rather than tethering the epidermal growth factor receptor to the basolateral membrane in *C. elegans*.

## Introduction

The spatial organization of signaling pathways is essential for proper activation and function of signal transduction cascades, particularly in the context of polarized cells (e.g. epithelial cells or neurons), in which cellular components are segregated into distinct domains. Mislocalization of signaling components can cause ectopic signaling activation or loss of signaling, leading to inappropriate cell responses in cell cycle regulation, migration, or survival that can lead to disease, developmental disorders, and cancer (Hung and Link, 2011).

Proper cellular organization is facilitated by adaptor and scaffolding proteins that ensure proximity of signaling components to activate signaling cascades, or that sequester components to inhibit activation (Buday and Tompa, 2010, Mugabo and Lim, 2018). For example, mammalian neurons rely on an evolutionarily conserved complex consisting of the adaptor and scaffolding proteins CASK (Calcium/calmodulin-dependent Serine protein Kinase), Lin7, and APBA1 (APP-Binding family A member 1) to maintain synaptic localization of the NMDA receptor subunit NR2B (Jeyifous et al., 2009, Jo et al., 1999, Setou et al., 2000), the synaptic adhesion molecule neurexin (Butz et al., 1998, Fairless et al., 2008), and the G-protein coupled receptor 5-HT2C (Becamel et al., 2002). Mutations in CASK, Lin7, and APBA1 are associated with cancer (Gruel et al., 2016, Hara et al., 2017, Wei et al., 2014), neurodegenerative diseases (Miller et al., 2006, Zucker et al., 2010), and cognitive dysfunction (Cristofoli et al., 2018, LaConte et al., 2014, Malik and Hodge, 2014), highlighting a role for these proteins in cellular function and human health. Lin7 and CASK are also expressed in epithelial cells, where they interact and coordinate polarized localization of the potassium channel Kir2.3 (Alewine et al., 2007). APBA1 has recently been found to be expressed in epithelial tissue (Motodate et al., 2016); however, whether it forms a complex with CASK and Lin7 in mammalian epithelia has not been shown.

An orthologous LIN-2 (CASK), LIN-7 (Lin7), and LIN-10 (APBA1) complex (the LIN-2/7/10 complex) exists in the genetic model organism *Caenorhabditis elegans,* where it is required for basolateral localization of the Epidermal Growth Factor Receptor (EGFR) homologue LET-23 in the progenitor epithelial cells that give rise to the vulva (Figure 1a) (Kaech et al., 1998). During the L2 and L3 larval stages of *C. elegans* development, basolateral LET-23 EGFR is necessary for activation of the downstream LET-60 Ras/MPK-1 ERK signaling cascade that specifies the primary vulval cell fate in the vulva precursor cells (VPCs). Of the six VPCs, only three cells are induced to generate the vulva due to their proximity to the source of the EGF-like ligand LIN-3. The remaining three VPCs assume the uninduced cell fate and divide once before fusing with the surrounding hypodermal syncytium, Hyp7. Loss-of-function mutations in *lin-2, lin-7,* or *lin-10,* or loss of interaction between LET-23 EGFR and LIN-7 results in exclusive apical localization of LET-23 EGFR, loss of signaling activation in the VPCs, and a vulvaless (Vul) phenotype (Aroian et al., 1994, Ferguson and Horvitz, 1985, Kaech et al., 1998). Increased LET-23 EGFR basolateral localization and signaling causes an excess of VPCs to assume vulval cell fates and leads to a multivulva (Muv) phenotype (Skorobogata et al., 2014).

**Figure 1:**
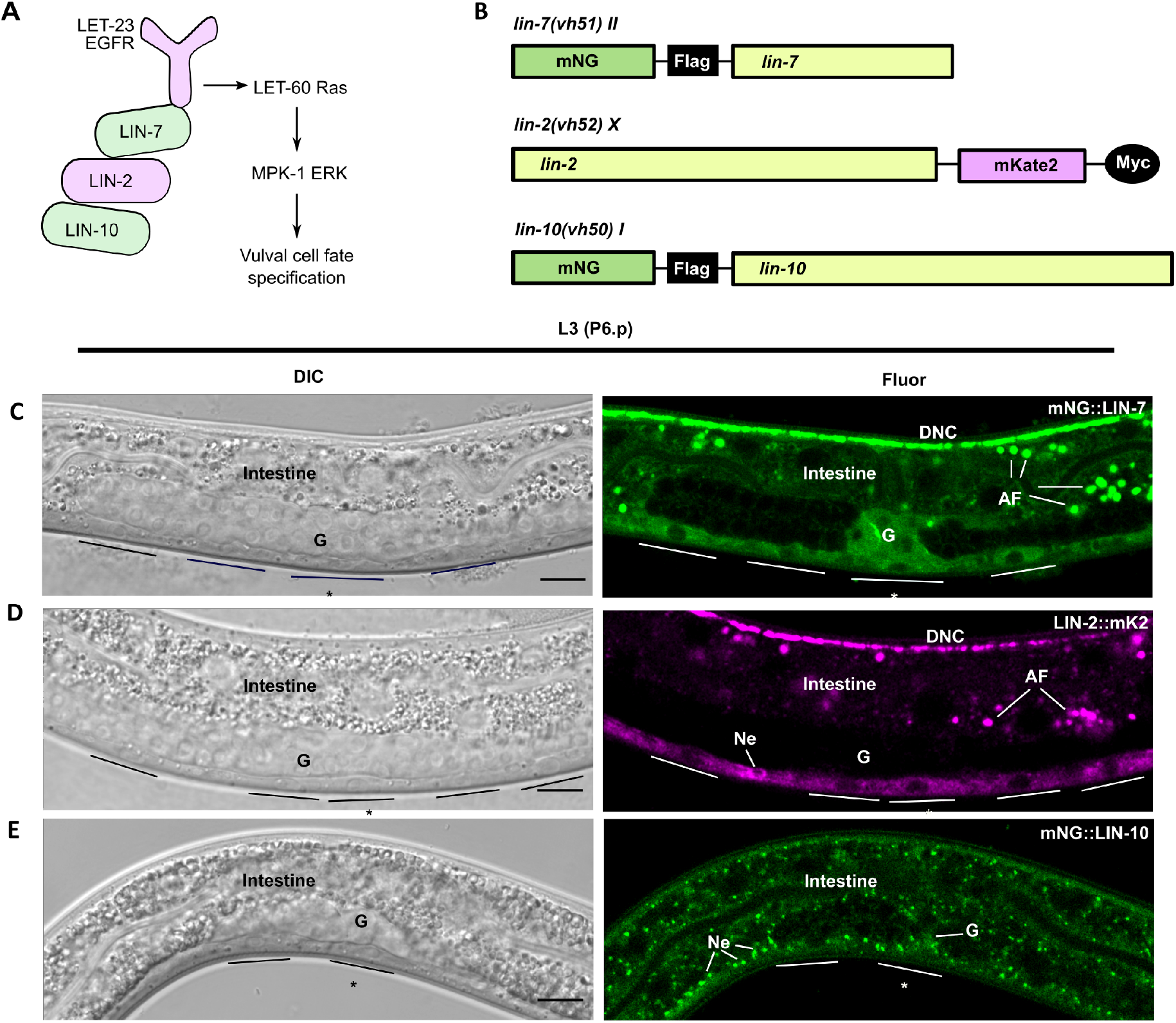
LIN-2, LIN-7, and LIN-10 are broadly expressed in *C. elegans*. **(a)** Schematic of how the LIN-2/7/10 complex interacts with the cytoplasmic tail of LET-23 EGFR in the vulva precursor cells, which is necessary for basolateral receptor localization and activation of the downstream Ras/ERK signalling cascade which specifies the vulval cell fate. **(b)** Schematic of endogenously-tagged *lin-7, lin-2,* and *lin-10* alleles generated by CRISPR/Cas9: *vh51, vh52,* and *vh50,* respectively. mNG: mNeonGreen. **(c-e)** Differential interference contrast (DIC) and corresponding confocal fluorescence images of L3 larvae (lateral view) expressing endogenously-tagged mNG::LIN-7 (c), LIN-2::mK2 (d) and mNG::LIN-10 (e). VPCs are underlined. Asterisk (*) denotes P6.p cell. G: Gonad. DNC: Dorsal nerve cord. AF: Non-specific autofluorescence in the intestine. Ne: Neuronal cell bodies in the ventral nerve cord. Scalebar: 10 μm.

The LIN-2/7/10 complex has been well-defined biochemically: *in vitro* interaction assays and yeast two-hybrid assays have shown that the C-terminal PDZ-interaction motif of LET-23 interacts with the C-terminal PDZ-domain of LIN-7, an interaction that is conserved with mammalian ErbB2 and ErbB4 (Kaech et al., 1998, Shelly et al., 2003, Simske et al., 1996). LIN-7 also interacts with LIN-2, which in turn interacts with LIN-10 (Kaech et al., 1998) (Figure 1a). These interactions have been validated in murine models (Borg et al., 1999, Borg et al., 1998, Butz et al., 1998) and the complex was co-immunoprecipitated from murine synaptic plasma membrane fractions (Butz et al., 1998).

In mammals, Lin7 and CASK colocalize on the basolateral membranes of epithelial cells (Cohen et al., 1998, Stetak et al., 2006, Straight et al., 2000), whereas CASK colocalizes with Golgi-associated APBA1 in neurons (Borg et al., 1999). Otherwise, CASK is typically found at presynaptic and postsynaptic termini (Chen and Featherstone, 2011, Hsueh, 2006). Individually, CASK and all three APBA paralogues have been shown to localize to the nucleus and regulate transcription (Hirose et al., 2014, Hsueh et al., 2000, Lau et al., 2010, Sumioka et al., 2008, Swistowski et al., 2009). Colocalization between Lin7 and APBA has not been demonstrated. The expression and subcellular localization of the LIN-2/7/10 complex in *C. elegans* has largely been unexplored. Overexpressed *C. elegans* LIN-7 roughly overlapped with apical junctions in the VPCs (Simske et al., 1996), whereas LIN-10 immunolocalized to cytoplasmic foci in descendants of the VPCs (Whitfield et al., 1999), suggesting that they may occupy distinct subcellular compartments. The localization of *C. elegans* LIN-2 has not been investigated in the VPCs. Identifying the subcellular localization of complex formation, and where the complex localizes with LET-23 EGFR, can provide insight into how the complex regulates polarized receptor localization, such as by tethering LET-23 EGFR at the basolateral membrane or through targeted secretion.

Towards gaining insight into how the LIN-2/7/10 complex coordinates membrane targeting of signaling receptors, we used CRISPR/Cas9 genome editing to tag endogenous LIN-2, LIN-7, and LIN-10 with fluorescent fusion proteins to identify their subcellular localization, dynamics, and expression in *C. elegans*. We found that LIN-7 is the only complex component that colocalizes with LET-23 EGFR on the basolateral membrane of the VPCs; however, this localization is independent of the LET-23 EGFR C-terminal tail previously shown to bind LIN-7. We identify intracellular compartments as the common site of localization of the LIN-2/7/10 complex and LET-23 EGFR, and that LIN-10 is largely required for recruitment of LIN-2 and LIN-7 to these compartments. We demonstrate that the LIN-2/7/10 complex forms *in vivo,* but that LIN-2/7 have different expression patterns from LIN-10, consistent with their stronger interactions. Our results suggest that the LIN-2/7/10 complex forms on Golgi ministacks or recycling endosomes to target LET-23 EGFR to the basolateral membrane of the VPCs rather than functioning as a tether at the basolateral membrane.

## Materials and Methods

### Strains and maintenance

*C. elegans* strains were maintained as previously described (Brenner, 1974). Worms were grown on *E. coli* HB101 at 20°C and all strains were derived from the N2 wild-type strain. A complete list of strains can be found in Table S1.

### Generating endogenously-tagged transgenic strains by CRISPR/Cas9

Primers used for cloning and genotyping are listed in Table S2. mNeonGreen::3xFlag::LIN-10 *(lin-10(vh50)),* mNeonGreen::3xFlag::LIN-7 *(lin-7(vh51)),* and LIN-2::3xMyc::mKate2 *(lin-2(vh52))* were generated by CRISPR/Cas9 cloning following the Self-Excising Cassette method (Dickinson et al., 2015). The following guide RNA sequences were used: 5’-ACCATGAACAATTCTGTTGC-3’ for *lin-10,* 5’-TTCCAGATGGATAACCCGGA-3’ for *lin-7,* and 5’-GATCAGTAGACCCAAGTGAC-3’ for *lin-2.* Guide RNAs were designed using a prediction software developed by Dr. Feng Zhang’s lab at the Massachusetts Institute of Technology (crispr.mit.edu).

### Generation of extrachromosomal array lines

Extrachromosomal *vhEx37* and *vhEx58* were cloned by inserting the open reading frames of *lin-10a* and *lin-2a* (amplified from wild-type cDNA, Table S2), respectively, downstream of codon-optimized GFP into a vector with a *lin-31* promoter and an *unc-54* 3’ untranslated region provided by Dr. Chris Rongo (Rutgers University, NJ). Extrachromosomal *vhEx63* was cloned by replacing the GFP of *vhEx37* with mCherry. Extrachromosomal *vhEx60* was cloned by inserting *lin-7a* open reading frame into a modified p255 vector expressing EGFP under a *lin-31* promoter (Skorobogata et al., 2014). All expression vectors (40 ng/μl) were injected with a *pttx-3::gfp* co-injection marker (80 ng/μl) following established protocols (Hobert et al., 1997, Mello et al., 1991).

### Microscopy and image analysis

Epifluorescent (black-and-white) images were acquired on a Zeiss Axio A1 Imager. Confocal (colorized) images were acquired using an LSM780 laser-scanning confocal microscope (Zeiss). Live animal imaging was performed as previously described (Sulston and Horvitz, 1977).

Punctate or membrane localization of LIN-2, −7, −10, and LET-23 were determined by visual inspection and confirmed by comparing fluorescence intensity of the region of interest to the background cytosolic fluorescence intensity. Peaks in fluorescence spanning the puncta or membrane region of at least twice the intensity of cytosolic fluorescence were categorized as punctate or membrane-associated.

Cytosolic fluorescence intensity was measured by sampling three 1 μm^2^ cytosolic regions (using ImageJ) in primary cell lineages free of any apparent punctae to achieve consistent measurements. Three 1 μm^2^ regions of background fluorescence were also randomly sampled from regions of the image with no worm present. The difference between the average cytosolic fluorescence intensity and the average background fluorescence was used as a measurement of fluorescence intensity for each image. At least 10 images were analyzed per vulval development stage (Figure 2a).

**Figure 2:**
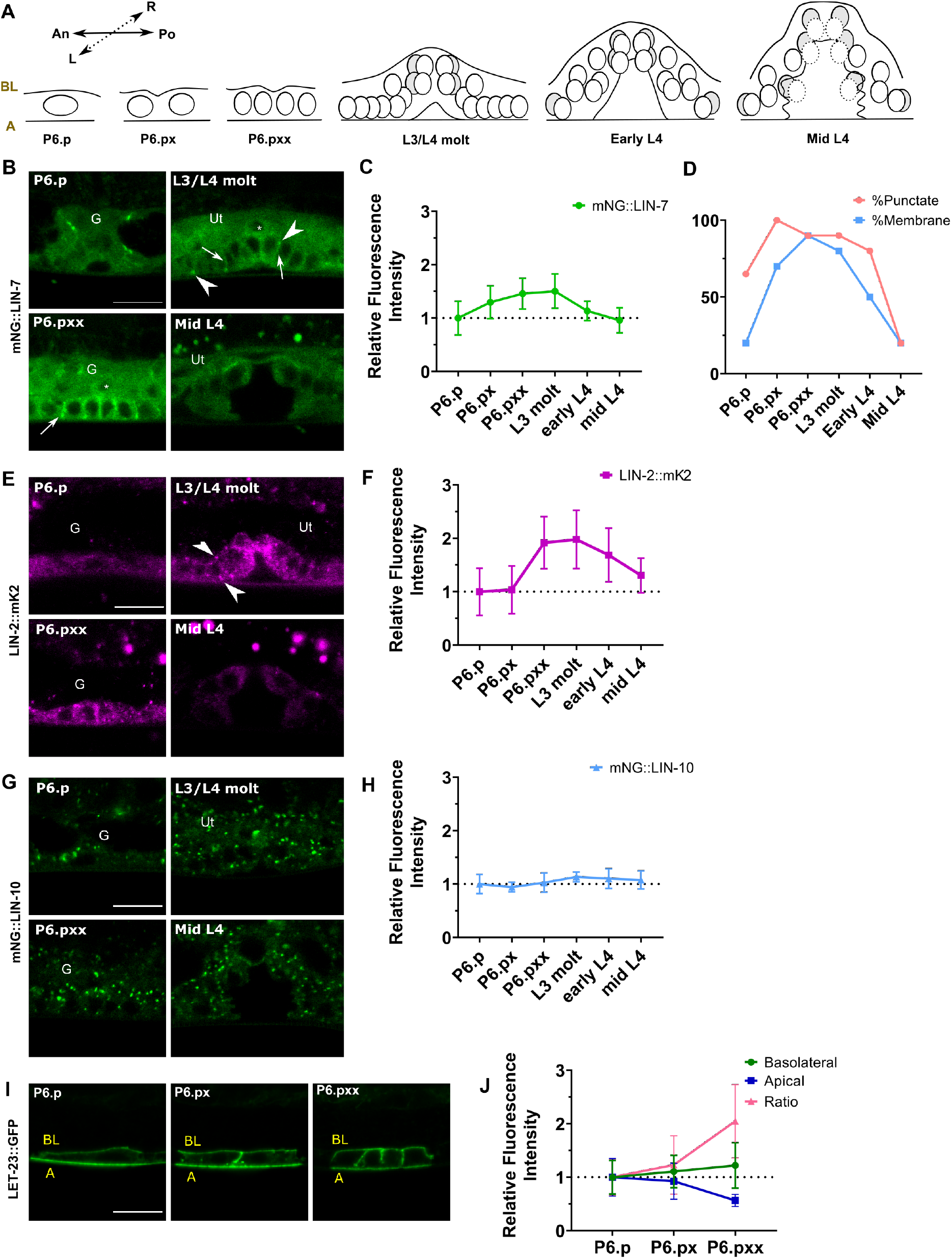
Expression and localization dynamics of LIN-2/7/10 and LET-23 EGFR. **(a)** Schematic of the stages of vulval development, from induction (late L2/early L3) to midmorphogenesis (mid L4), used for analysis of fluorescent intensity. An, Anterior. Po, Posterior. L, Left. R, Right. **(b)** mNG::LIN-7 expression and localization in P6.p, P6.pxx, L3/L4 molt, and mid L4 worms. Arrowhead: punctate localization of LIN-7. Arrow: membrane localization of LIN-7. Asterisk: nucleus of anchor cell. **(c)** mNG::LIN-7 cytosolic fluorescent intensity expression analysis from P6.p to mid-L4. **(d)** Analysis of mNG::LIN-7 localization patterns from P6.p to mid-L4. **(e)** LIN-2::mK2 expression and localization in P6.p, P6.pxx, L3/L4 molt, and mid L4 worms. Arrowhead: punctate localization of LIN-2. **(f)** LIN-2::mK2 cytosolic fluorescent intensity analysis from P6.p to mid-L4. **(g)** mNG::LIN-10 expression and localization in P6.p, P6.pxx, L3/L4 molt, and mid L4 worms. **(h)** mNG::LIN-10 cytosolic fluorescent intensity analysis from P6.p to mid-L4. **(i)** LET-23::GFP *(zhIs035)* expression and localization in P6.p, P6.px, and P6.pxx worms. **(j)** Peak basolateral, apical, and basolateral/apical ratio fluorescent intensity analysis of LET-23::GFP. BL: Basolateral. A: Apical. G: Gonad. Ut: Uterus. Scalebar: 5 μm.

Polarized membrane distribution of LET-23 EGFR was analyzed as previously described by drawing a 20 pixel-wide line across P6.p cell nuclei for each worm and measuring the peak fluorescence intensity on the basolateral and apical membrane (Skorobogata et al., 2014).

Mander’s correlation coefficients for endogenously-tagged LIN-2, LIN-7, LIN-10, and LET-23 EGFR were measured using Zen 2012 software (Zeiss) and quantified by tracing the cells of interest and setting a crosshair threshold that omits background fluorescence and includes cytosolic signal. The same thresholds were used for all images analyzed. Overlap of punctae was determined by identifying punctae using methods described above and checking for overlap in peak fluorescence intensity. Average number of overlapping punctae per worm was used for analysis.

### Analyzing VPC cell fate induction

VPC induction scoring was performed as described previously (Gauthier and Rocheleau, 2017). Each worm was given a VPC induction score between 0 and 6, with scores less than 3 classified as Vul and greater than 3 as Muv. For vulval induction of worms expressing extrachromosomal arrays, worms expressing GFP in any VPC lineage were scored and compared to siblings on the same plate lacking any detectable extrachromosomal array expression (including the co-injection marker, *pttx-3::gfp).*

### Co-immunoprecipitation

Six NGM plates saturated with healthy worms kept at 20°C were washed off using sterile M9 buffer and collected in a 15 ml conical tube for each genotype. Harvested worms were washed three times with fresh M9, and two additional times with chilled (4°C) worm protein lysis buffer (50 mM Hepes pH 7.6, 1 mM EDTA, 1 mM MgCl_2_, 100 mM KCl, 10% glycerol, 0.05% NP-40, cOmplete™ EDTA-free Protease Inhibitor Cocktail tablet [Sigma], NaF, Na_3_VO_4_, PMSF). Worm pellets were resuspended in fresh lysis buffer (up to 2 ml) and freeze/thawed five times in liquid nitrogen, sonicated three times (30% amplitude, 3 s on and 5 s off for a total sonication time of 15 seconds) until the sample was homogenized, and centrifuged at 12,000 x g for 30 min (4°C). Supernatant was collected as the worm lysate. SureBeads™ Protein G Magnetic Beads (BioRad) were washed with 100 μl worm protein lysis buffer and incubated with monoclonal M2 mouse anti-Flag antibody (Sigma, F3165) or monoclonal rabbit anti-c-Myc antibody (Sigma, PLA0001) in lysis buffer for 1 h at room temperature while rotating on a nutator. Antibody-bound beads were incubated with 800 μg whole worm lysate overnight at 4°C while rotating. Protein concentration was measured using BSA standard assay with Bradford reagent (BioRad). Precipitated proteins were eluted from the beads by boiling in 30 μl 1X SDS sample buffer for 10 min prior to electrophoresis. Co-immunoprecipitation assay was performed four times for each condition.

SDS-PAGE was performed using TGX Stain-Free FastCast mini gels (BioRad). Protein content on gels were then transferred onto PVDF membranes. Membranes were blocked for 1 h with 5% skim milk in 0.1% TBS-T and probed with 1:2000 primary antibody for bait and prey proteins (mouse anti-Flag for LIN-10 or LIN-7, rabbit anti-c-Myc for LIN-2) diluted in blocking solution and incubated overnight at 4°C while rotating. The next day, membranes were washed with 0.1% TBS-T and incubated with 1:10,000 secondary antibody (rabbit anti-mouse or goat anti-rabbit [Sigma]), diluted in blocking solution for 1 h at room temperature. Membranes were exposed using ECL-Clarity (BioRad) and imaged using a ChemiDoc Imager (BioRad). For bait and prey of a similar size (LIN-10 and LIN-2), the membrane was stripped using a mild stripping buffer (Harlow and Lane, 2006), blocked for 1 h, and re-probed. In these cases, prey protein was probed first, then bait protein was probed after stripping.

### Statistical analysis

All statistical analyses were performed using GraphPad Prism 8.0. Two-tailed Student’s *t-*test or One-Way ANOVA with Dunnett’s test for multiple comparisons were used to compare average means. Fisher’s exact test was used to compare vulvaless phenotypes (Vul vs not-Vul), multivulva phenotypes (Muv vs not-Muv), and localization analyses (e.g. punctate vs not punctate).

## Results

### Localization and expression of LIN-2/7/10 in *C. elegans* VPCs

To identify where the LIN-2/7/10 complex forms *in vivo,* we used CRISPR/Cas9 to tag the endogenous 5’ end of *lin-7* and *lin-10* gene loci with an mNeonGreen (mNG) fluorophore and a 3xFlag tag, generating the *lin-7(vh51)* (“mNG::LIN-7”) and *lin-10(vh50)* (“mNG::LIN-10”) alleles, respectively. We also inserted an mKate2 (mK2) fluorophore and a 3xMyc tag to the 3’ end of the *lin-2* gene loci, generating the *lin-2(vh52)* (“LIN-2::mK2”) allele (Figure 1b). The gene products are predicted to generate wild-type, functional proteins based on the absence of vulval development defects in their respective lines (Table S3). For tissue-specific expression, we generated extrachromosomal array transgenes under a VPC-specific promoter *(lin-31)* of the LIN-7a isoform tagged C-terminally with EGFP *(vhEx60,* “LIN-7a::EGFP”), LIN-2a tagged N-terminally with GFP *(vhEx58,* “GFP::LIN-2a”), and LIN-10a tagged N-terminally with GFP *(vhEx37,* “GFP::LIN-10a”) (Figure S1a). These transgenes rescued the vulvaless phenotypes of their respective mutants, confirming functionality (Table S4).

In the VPCs, endogenous mNG::LIN-7 was strongly cytosolic (Figure 1c, 2b). LIN-7 frequently localized to both basolateral membranes and cytoplasmic foci in both L3 and L4 larval stage worms (Figure 2b and d, 8a-c). C-terminally-tagged extrachromosomal LIN-7a::EGFP, on the other hand, is strongly nuclear and cytosolic with no detectable punctate or membrane localization (Figure S1b). The C-terminal PDZ domain has previously been found to be required for cell junction localization of *C. elegans* LIN-7 and mammalian Lin7 in epithelia (Simske et al., 1996, Straight et al., 2000); therefore, placement of the fluorophore at the C-terminus might disrupt recruitment to membranes without compromising overall function. Alternatively, overexpression of LIN-7 from the extrachromosomal array may overwhelm any detectable signal at the plasma membrane or cytosolic foci.

Endogenous LIN-2::mK2 was also found to have a strong cytosolic signal and to localize to cytoplasmic foci in almost all L3 and L4 larvae, but did not have a distinct membrane localization pattern (Figure 1d, 2e, 8e-f). N-terminally tagged extrachromosomal GFP::LIN-2a was strongly cytoplasmic and nuclear. It also localized to punctae, although less frequently: only 30% of VPCs imaged had punctate LIN-2 localization (Figure S1c, 2).

Finally, both endogenous mNG::LIN-10 and extrachromosomal GFP::LIN-10 strongly localized to punctae with a relatively low cytosolic signal (Figure 1e, 2g, Figure S1d), consistent with localization to Golgi ministacks and recycling endosomes previously identified in neuronal and intestinal cells (Glodowski et al., 2005, Rongo et al., 1998, Zhang et al., 2012). Similar punctate localization for endogenous untagged LIN-10 has been found in immunostaining assays at the P6.pxx four-cell stage (Whitfield et al., 1999).

### Expression of LIN-2, LIN-7, and LET-23 EGFR, but not LIN-10, change throughout vulval development

We examined expression patterns for LIN-2/7/10 complex components throughout vulval development (Figure 2a) and found that cytosolic fluorescence intensities associated with LIN-2::mK2 and mNG::LIN-7 increase gradually from the one-cell P6.p stage and peak after all cell divisions have taken place near the L3/L4 molt, then drop in L4 larvae (Figure 2b-c, e-f). Furthermore, their expression is restricted to the induced vulval cell fate lineages: in the uninduced P3.p, P4.p, and P8.p cells, and in the uninduced cells of *lin-2(e1309)* or *lin-7(e1413)* null mutants, fluorescence intensity of LIN-2::mK2 and mNG::LIN-7 drops after one cell division, potentially coincident with cell fusion with Hyp7 (Figure 3a-d). In addition to changes in fluorescence intensity, mNG::LIN-7 displays changes in membrane and punctate localization throughout VPC induction. The proportion of worms with distinct mNG::LIN-7 membrane localization increases from 20% of one-cell P6.p to 90% of four-cell P6.pxx, then drops back to 20% in the developing vulva of mid-L4 larvae (Figure 2d). While most animals had some faint LIN-7-positive punctae throughout most of vulval development, the proportion of worms with punctate LIN-7 localization sharply decreases to 20% in mid-L4 vulva. On the other hand, LIN-2 and LIN-10 localization patterns are relatively unchanged throughout vulval development.

**Figure 3:**
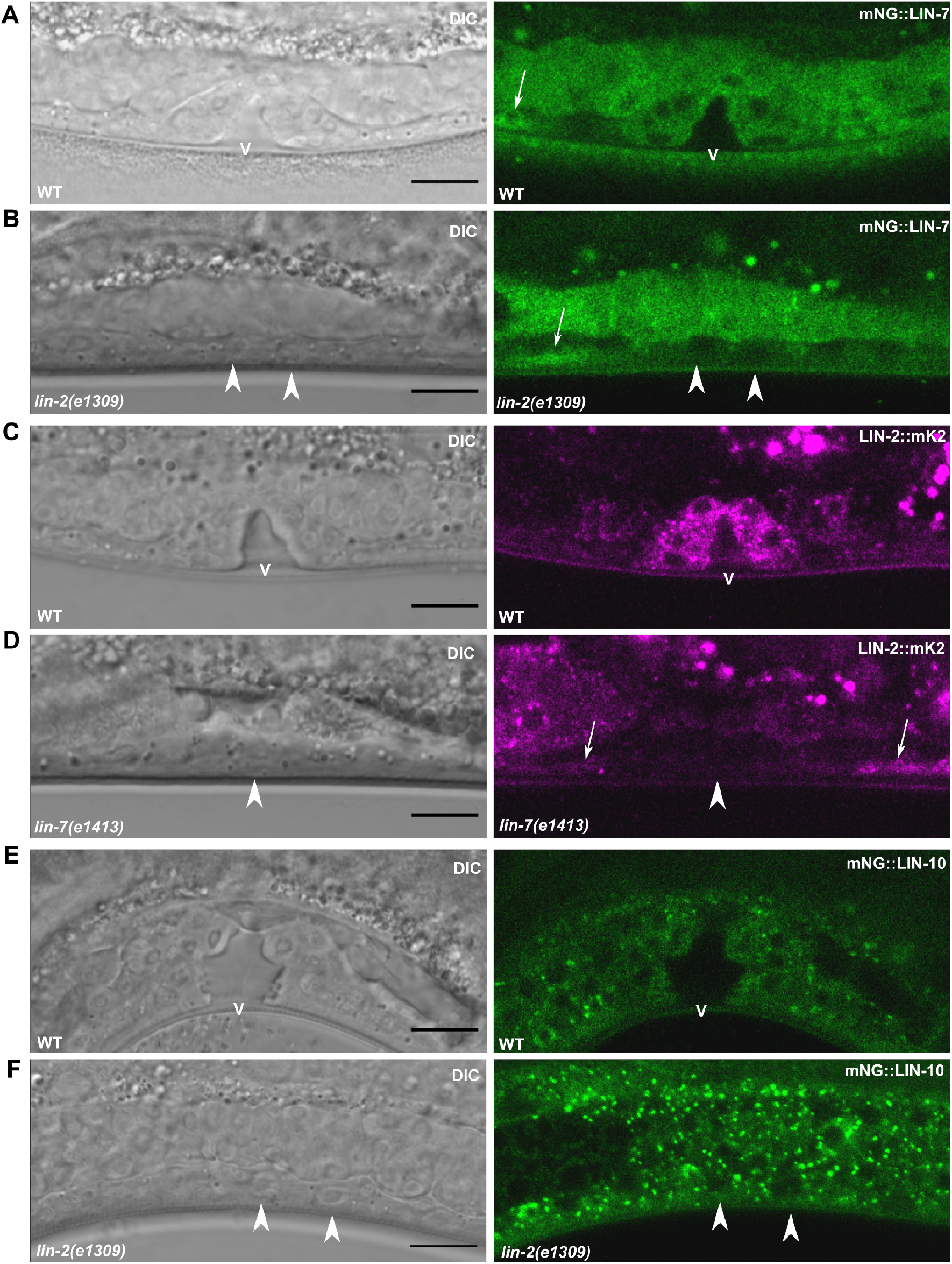
Expression of LIN-2 and LIN-7 is restricted to induced vulval cells. **(a-b)** mNG::LIN-7 expression in VPC lineages of wildtype (a) and *lin-2* mutant (b) L4 larvae. **(c-d)** LIN-2::mK2 expression in VPC lineages of wildtype (c) and *lin-7* mutant (d) L4 larvae. **(e-f)** mNG::LIN-10 expression in VPC lineages of wildtype (e) and *lin-2* mutant (f) L4 larvae. Scalebars: 5 μm. Arrowhead: nuclei of uninduced cells. Arrow: segment of ventral nerve chord in same focal plane as the VPCs. V: Vulval lumen.

Unlike its complex components, LIN-10 cytosolic fluorescence intensity does not detectably change throughout vulval development. LIN-10 is expressed evenly in all VPCs and their descendants, including the non-vulval cell lineages of P3.p, P4.p, and P8.p (Figures 2g-h, and 3e-f).

Membrane-bound LET-23 EGFR is localized in a polarized fashion in the VPCs, with stronger fluorescence intensity detected on the apical membrane than the basolateral membrane (Skorobogata et al., 2014). To look for changes in LET-23 EGFR localization, we compared the peak fluorescence intensities of an integrated LET-23::GFP transgene *(zhIs035* (Haag et al., 2014)) along the basolateral and apical membranes of P6.p, P6.px, and P6.pxx cells. The subsequent cell division generating P6.pxxx is a transverse division along the left/right plane and occurs with the formation of the apical lumen of the vulva (Sternberg and Horvitz, 1986) (Figure 2a), and as a result the apical membranes of P6.p lineages face the lumen rather than the ventral side (Sharma-Kishore et al., 1999), obscuring them from imaging and analysis. We found that the basolateral/apical fluorescence intensity ratio of LET-23::GFP doubles from P6.p to P6.pxx, coinciding with a drop in fluorescence intensity on the apical membrane (Figure 2i-j).

### LIN-2 and LIN-7 colocalize strongly with each other, and occasionally with LIN-10 at cytoplasmic foci in the VPCs

To identify where the LIN-2/7/10 complex forms *in vivo,* we crossed both *lin-7(vh51)* and *lin-10(vh50)* with *lin-2(vh52)*. We found that interacting partners LIN-7 and LIN-2 colocalize strongly in the cytosol of the VPCs and at cytoplasmic foci (Figure 4a), although LIN-2 is frequently localized to punctae without LIN-7. Mander’s correlation coefficients reveal moderately strong colocalization between LIN-2 and LIN-7 (Figure 4c-d).

**Figure 4:**
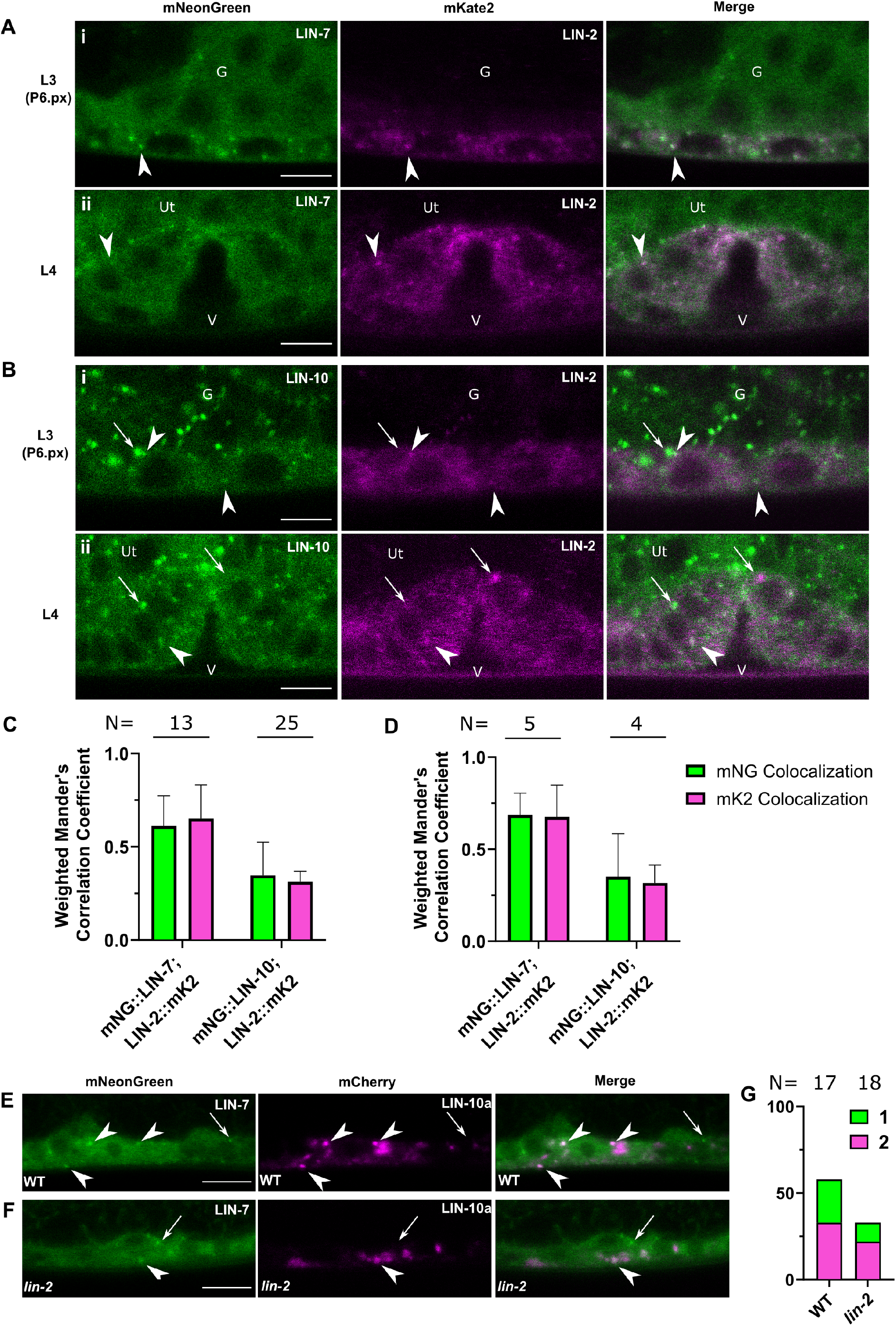
The LIN-2/7/10 complex colocalizes on cytoplasmic punctae. **(a)** mNG::LIN-7 and LIN-2::mK2 colocalize in the cytosol and at punctae in L3 (i) and L4 (ii) worms. **(b)** mNG::LIN-10 and LIN-2::mK2 colocalize at some punctae in L3 (i) and L4 (ii) worms. **(c-d)** Weighted Mander’s colocalization coefficients for L3 (c) and L4 (d) larval stages. **(e-f)** Overlap of mNG::LIN-7-positive with mCherry::LIN-10a punctae in a wildtype (e) and *lin-2* mutant (f) background. **(g)** Quantification of the percentage (on Y-axis) of VPCs imaged with punctate mNG::LIN-7 localization (1, green), and the percentage of LIN-7-positive punctae that overlaps with mCh::LIN-10a (2, magenta) in the wild type and *lin-2* mutants from (e-f). Scalebars: 5 μm. Arrowhead: colocalizing punctae. Arrow: noncolocalizing punctae. V: Vulval lumen. G: L3 gonad. Ut: L4 uterus. Error bars: SD.

There is some overlap between LIN-2 and LIN-10 at cytoplasmic punctae; however, they are frequently localized to separate compartments (Figure 4b). Mander’s colocalization coefficients reveal relatively weak colocalization between LIN-10 and LIN-2 (Figure 4c-d).

To test for overlap between LIN-7 and LIN-10, we crossed *lin-7(vh51)* with an extrachromosomal mCherry-tagged LIN-10a transgenic line *(vhEx63)* (mCh::LIN-10), and found that approximately 60% of LIN-7-positive punctae overlap with LIN-10-positive punctae. The overlap between LIN-7 and LIN-10 is in part LIN-2-dependent, as LIN-7 localization to punctae decreases in a *lin-2(e1309)* mutant (Figure 4e-g), described further below (Figure 8). The small number of LIN-7-positive punctae present in a *lin-2* mutant overlap with mCh::LIN-10 at a similar frequency as a wildtype background, suggesting that a small number of LIN-7-positive punctae localizes to similar subcellular compartments as LIN-10, even without LIN-2 (Figure 4e-g).

### LET-23 EGFR colocalizes with LIN-7 at the plasma membrane and with LIN-10 at cytoplasmic foci in the VPCs

To identify where the LIN-2/7/10 complex might interact with LET-23 EGFR, we crossed *lin-7(vh51)* and *lin-10(vh50)* with a strain expressing endogenously-tagged LET-23::mKate2::3xFlag *(let-23(re202);* “LET-23::mK2”; generated by CRISPR/Cas9; generously provided by T. Duong and D. Reiner). LET-23::mK2 localizes to the basolateral and apical membrane domains of the VPCs (Figure 5a-b), as has been described for endogenous LET-23 EGFR and other LET-23 EGFR reporters (Haag et al., 2014, Skorobogata et al., 2014, Whitfield et al., 1999). The fluorophore is placed upstream of the PDZ interaction motif on the C-terminal end to preserve the interaction with LIN-7, similar to other functional LET-23::GFP transgenes (Haag et al., 2014). However, basolateral receptor localization is at times undetectable at the one-cell P6.p stage, and the presence of mild and infrequent vulval abnormalities suggests the modifications made to the endogenous *let-23* gene locus cause a minor disruption to its regular function (Table S3).

**Figure 5:**
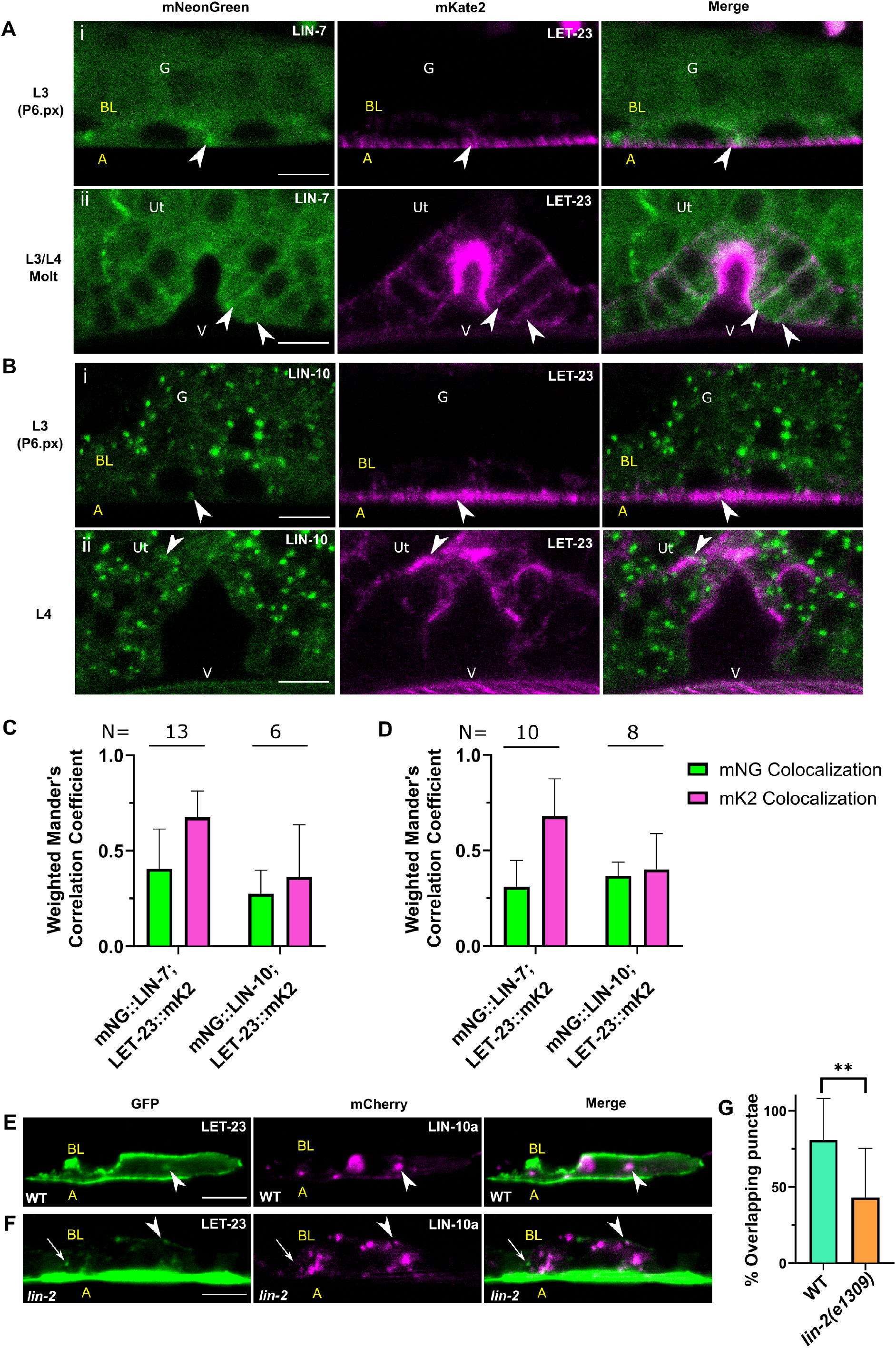
LET-23 EGFR colocalizes with LIN-7 at basolateral membranes and with LIN-10 at foci. **(a)** mNG::LIN-7 and LET-23::mK2 *(re202)* overlap at basolateral membranes in L3 (i) and L4 (ii) worms. **(b)** mNG::LIN-10 and LET-23::mK2 infrequently overlap in L3 (i) and L4 (ii) worms. **(c-d)** Weighted Mander’s colocalization coefficients for L3 (c) and L4 (d) larval stages. **(e-f)** Overlap of faint LET-23::GFP punctae with mCherry::LIN-10a-positive punctae in wildtype (e) and a *lin-2* mutant (f). **(g)** Quantification of LET-23::GFP-positive punctae that overlap with mCh::LIN-10a-positive punctae, as shown in (e-f). N = 10 P6.p cells for WT, 11 for *lin-2(e1309).* **p<0.01 Two-tailed Student’s *t*-test. Scalebars: 5 μm. Arrowhead: colocalizing punctae/membranes. Arrow: non-colocalizing punctae. V: Vulval lumen. G: L3 gonad. Ut: L4 uterus. A: Apical. BL: Basolateral. Error bars: SD.

We found that LET-23::mK2 and mNG::LIN-7 overlap at the basolateral plasma membrane (Figure 5a) in the VPCs (L3) and differentiated vulval cells (L4). This suggests that LIN-7 interacts with the receptor at the cell periphery. This interaction might happen in the absence of LIN-2 because LIN-2::mK2 does not reveal any apparent membrane localization (Figure 4a). Mander’s correlation coefficients show that mNG::LIN-7 colocalizes weakly with LET-23::mK2; however, the receptor colocalizes with mNG::LIN-7 relatively strongly (Figure 5c-d).

There was minimal colocalization between LET-23::mK2 and mNG::LIN-10 (Fig 5b-d), although LIN-10 does occasionally overlap with LET-23 EGFR at the cell periphery. We previously found that LET-23::GFP typically localizes to a few small cytoplasmic foci in most P6.p and P6.px cells (Skorobogata et al., 2016). LET-23::mK2 fluorescence intensity was too low to detect cytosolic punctae; therefore, to determine if LIN-10 might colocalize with LET-23 EGFR intracellularly, we crossed the LET-23::GFP integrated transgene with extrachromosomal mCh::LIN-10a and found some overlap between cytosolic LET-23::GFP-positive foci with LIN-10 (Figure 5e). This overlap is significantly decreased in a *lin-2* mutant (Figure 5f-g).

### LIN-2 and LIN-7 colocalize in neurons, but not with LIN-10 and LET-23

Endogenously-tagged LIN-2, LIN-7, LIN-10, and LET-23 EGFR allow for analysis of their localization and expression patterns in other tissues. We found that all four proteins are expressed in neurons and sensory tissue in the anterior half of the worm. The intestine is prone to a high degree of autofluorescence, and is excluded from this initial analysis. Whereas LET-23 EGFR and LIN-7 overlap minimally in the head (Figure 6a), LIN-2 and LIN-7 colocalize strongly in the neural ring, and the ventral and dorsal nerve chords (Figure 6b). Of note, LIN-2 is more strongly expressed in the isthmus of the pharynx than LIN-7 (Figure 6b). In contrast LIN-7 is more strongly expressed in the gonad and uterus than LIN-2 (see Figures 1–5). LIN-10 overlaps minimally with LIN-2 in the neural ring and nerve chords (Figure 6c), and shares very little overlap with LET-23 EGFR in other neural tissues in the head of the worm (Figure 6d). LET-23::mK2 is strongly expressed in the excretory duct cell (Figure 6a, d) where it signals through the LET-60 Ras/MPK-1 ERK pathway to regulate excretory duct cell development (Abdus-Saboor et al., 2011, Yochem et al., 1997). Loss-of-function mutations in *let-23,* but not in *lin-2/7/10* complex components, yield a larval lethal phenotype associated with loss of the excretory duct cell (Ferguson and Horvitz, 1985). The LIN-2/7/10 complex components were not observed in the excretory duct cell. Further analysis is needed to identify the precise cells in which LIN-2, LIN-7, and LIN-10 are expressed.

**Figure 6:**
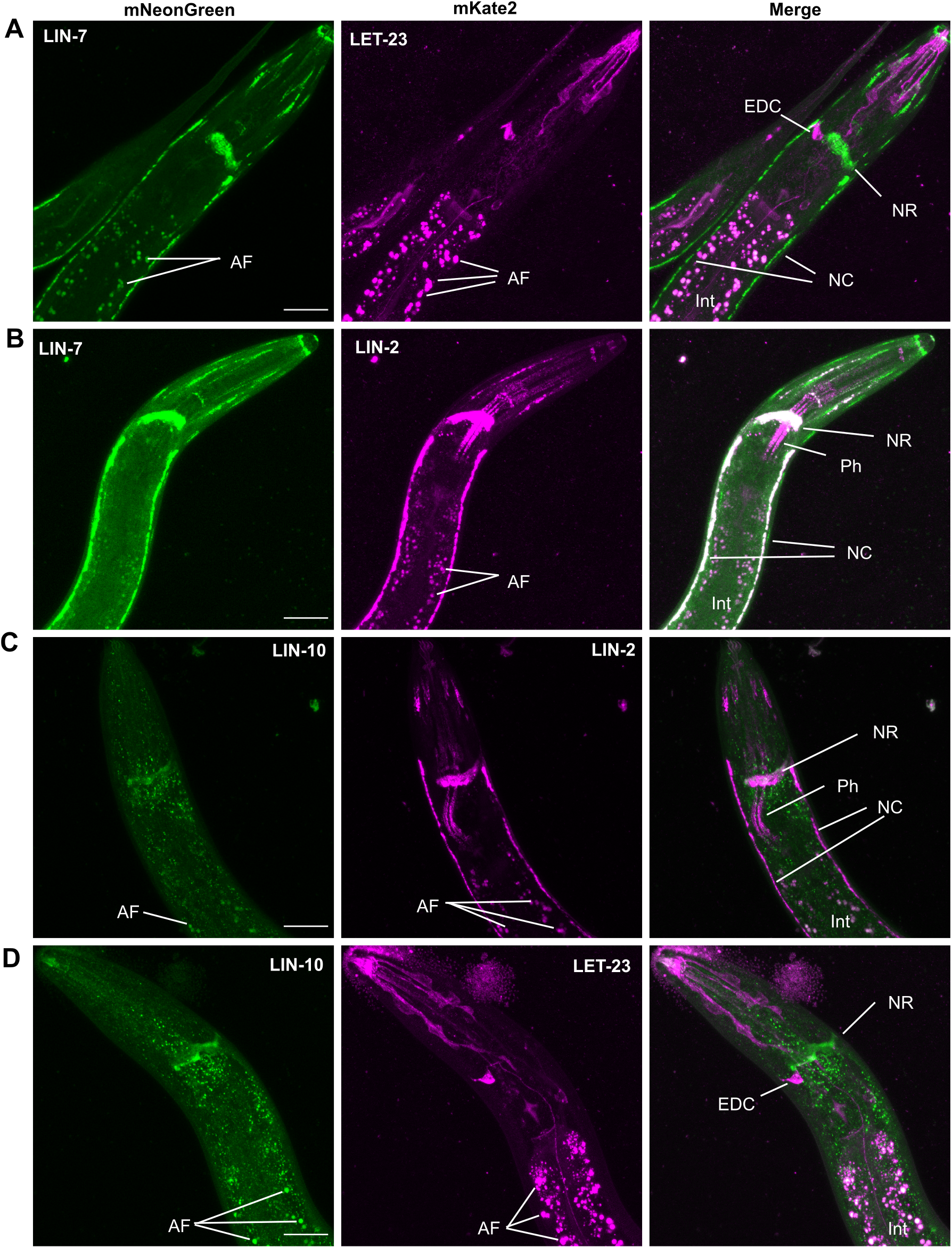
LIN-2 and LIN-7, but not LIN-10 or LET-23 EGFR, colocalize in neurons. **(a-d)** Three-dimensional Z-stack maximum intensity projections of the anterior half of L3 (ac) and L4 (d) larval worms. **(a)** mNG::LIN-7 and LET-23::mK2 EGFR were both expressed in several tissues in the head of *C. elegans* but do not colocalize in overlap. **(b)** mNG::LIN-7 and LIN-2::mK2 colocalized strongly in neuronal tissues. **(c)** LIN-2::mK2 and LIN-10::mNG overlapped minimally in nerve ring. **(d)** LIN-10::mNG and LET-23::mK2 EGFR had distinct localization and expression patterns in the head. Scalebars: 20 μm. AF: Autofluorescence. EDC: Excretory Duct Cell. Int: Intestine. NC: Nerve Chords. NR: Neural Ring. Ph: Pharynx.

### LIN-2::mK2 interacts strongly with mNG::LIN-7 and minimally with mNG::LIN-10 *in vivo*

The specific interactions between LIN-2 and LIN-7, and between LIN-2 and LIN-10 have been tested by yeast two-hybrid assay, and complex formation has been confirmed by coimmunoprecipitation of the *C. elegans* proteins expressed exogenously in Drosophila S2 cells (Kaech et al., 1998). These interactions have been shown to be evolutionarily conserved, and the complex has been shown to form in mammalian neurons by co-immunoprecipitation and *in vitro* pull-down assays (Becamel et al., 2002, Borg et al., 1998, Butz et al., 1998, Leonoudakis et al., 2004). It has not yet been shown that the complex forms *in vivo* in *C. elegans*, and the disparity in localization patterns observed in Figures 4 and 6 call into question the extent of these interactions *in vivo.*

We performed a co-immunoprecipitation assay using whole worm lysates to test if the proteins interact *in vivo* (Figure 7). On a western blot, we found that mNG::3xFlag::LIN-10 yielded two bands. The larger band at roughly 180 kDa is expected to include the three known isoforms of LIN-10 (a, b, and c), and is about 45 kDa larger than anticipated of the fusion of LIN-10 with a fluorophore. A similar size shift has been previously reported using antibodies for endogenous LIN-10, and could be due to extensive post-translational modifications (Whitfield et al., 1999). A second smaller band was also observed for LIN-10 as expected from published work, and may represent an uncharacterized splice variant or proteolytic cleavage (Whitfield et al., 1999). Blotting for mNG::3xFlag::LIN-7 yielded the expected single band at around 65 kDa, and LIN-2::mK::3xMyc yielded the expected two bands at 140 kDa and 100 kDa representing the full-length a isoform and shorter b isoform, respectively (Figure 7).

**Figure 7:**
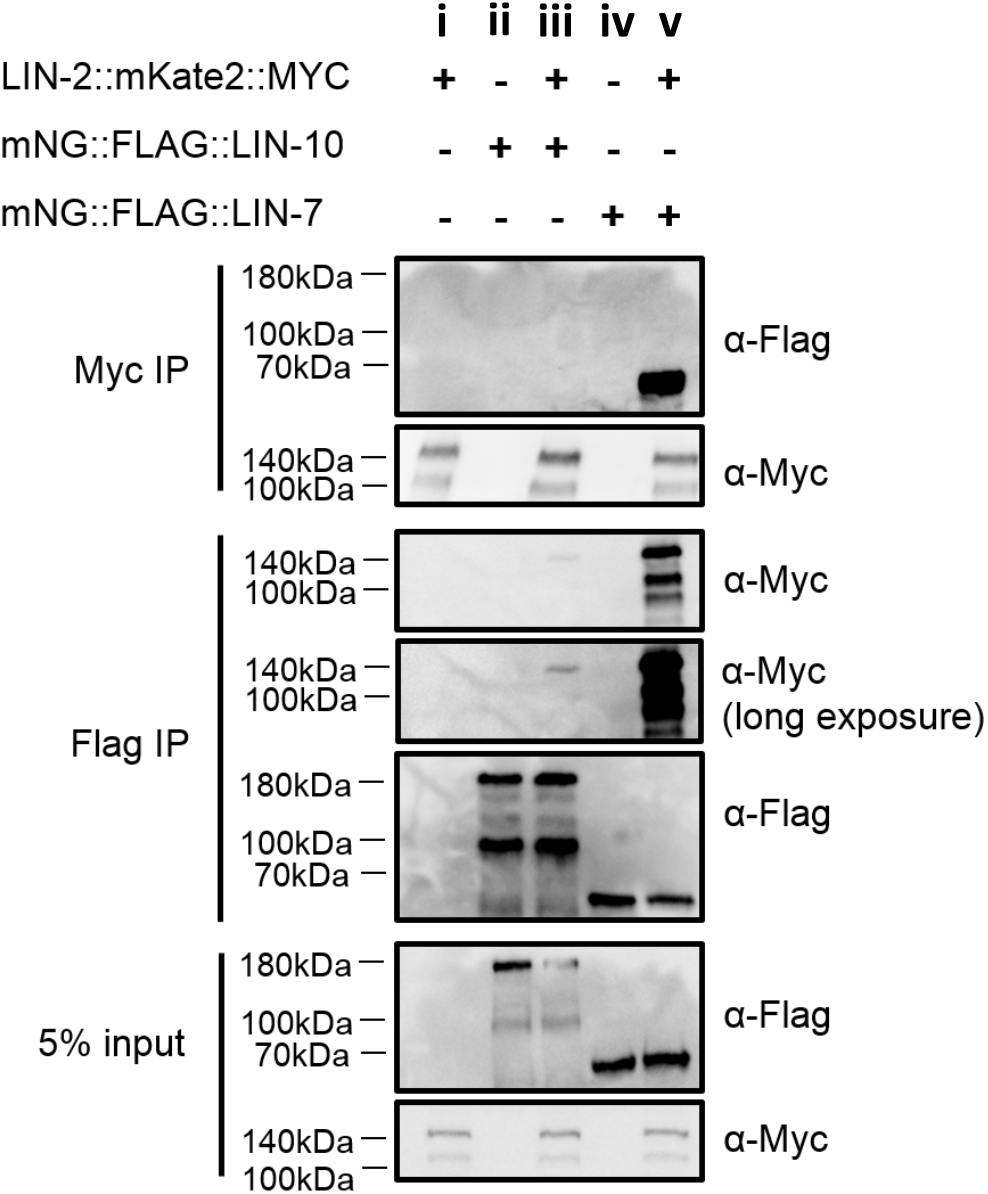
LIN-2, LIN-7, and LIN-10 interact *in vivo*. Co-immunoprecipitation assays using whole worm lysate from worms expressing: (i) LIN-2::mK2::3xMyc, (ii) mNG::3xFlag::LIN-10, (iii) both LIN-2::mK::3xMyc and mNG::3xFlag::LIN-10, (iv) mNG::3xFlag::LIN-7, and (v) both LIN-2::mK::3xMyc and mNG::3xFlag::LIN-7.

The full-length LIN-2a isoform co-immunoprecipitated with both LIN-7 and LIN-10 from whole animal lysates. On the other hand, the truncated LIN-2b isoform lacking the N-terminal CamKII domain only precipitated with LIN-7, consistent with this domain being required for interaction with LIN-10. Furthermore, while LIN-7 and LIN-2 coimmunoprecipitated strongly together, LIN-2 co-immunoprecipitated weakly with LIN-10, and no LIN-10 was recovered when purifying LIN-2 from lysates (Figure 7). These results and the colocalization analysis suggest LIN-2 and LIN-7 interact stably in several tissues, whereas LIN-2 and LIN-10 interact minimally.

### LIN-10 recruits LIN-2 and LIN-7 to subset of cytosolic punctae

To further characterize the localization dynamics of LIN-2, LIN-7, and LIN-10, we tested their interdependency for localization. Given the strong colocalization between LIN-2 and LIN-7, and their dependency for localization in some mammalian epithelial cells (Lozovatsky et al., 2009, Straight et al., 2000), we reasoned that they may rely on each other for localization in the VPCs as well. Because mNG::LIN-7 and LIN-2::mK2 expression was lost in uninduced cells (Figure 3), we limited our analysis to P6.p cells in L3 worms (before cell fate determination), and to L4 worms with partial or full vulval development (in which LIN-7 and LIN-2 are expressed). We found that LIN-7 punctate localization is largely LIN-2-and LIN-10-dependent. mNG::LIN-7, which localizes to punctae in 65% of L3 worms (P6.p) and 50% of all L4 worms (pooled for both early- and mid-L4), was localized to punctae in only 15% of P6.p cells and 10% of L4 worms with partial or full vulval development in *lin-2(e1309)* null mutants (Figure 8a-c). In *lin-10(e1439)* null mutants, punctate localization of LIN-7 was similarly decreased to 15% of P6.p cells and 15% of L4 worms with partial or full vulval development (Figure 8a-c). LIN-7 basolateral membrane localization, on the other hand, is slightly not but significantly decreased in *lin-2(e1309)* mutants (Figure 8b-c). Membrane localization of LIN-7 appeared to be LIN-10-dependent in P6.p cells, where its association with the basolateral membrane decreased from 20% in wildtype to 0% in *lin-10(e1439)* mutants, although this was not found to be statistically significant (Figure 8b-c).

**Figure 8:**
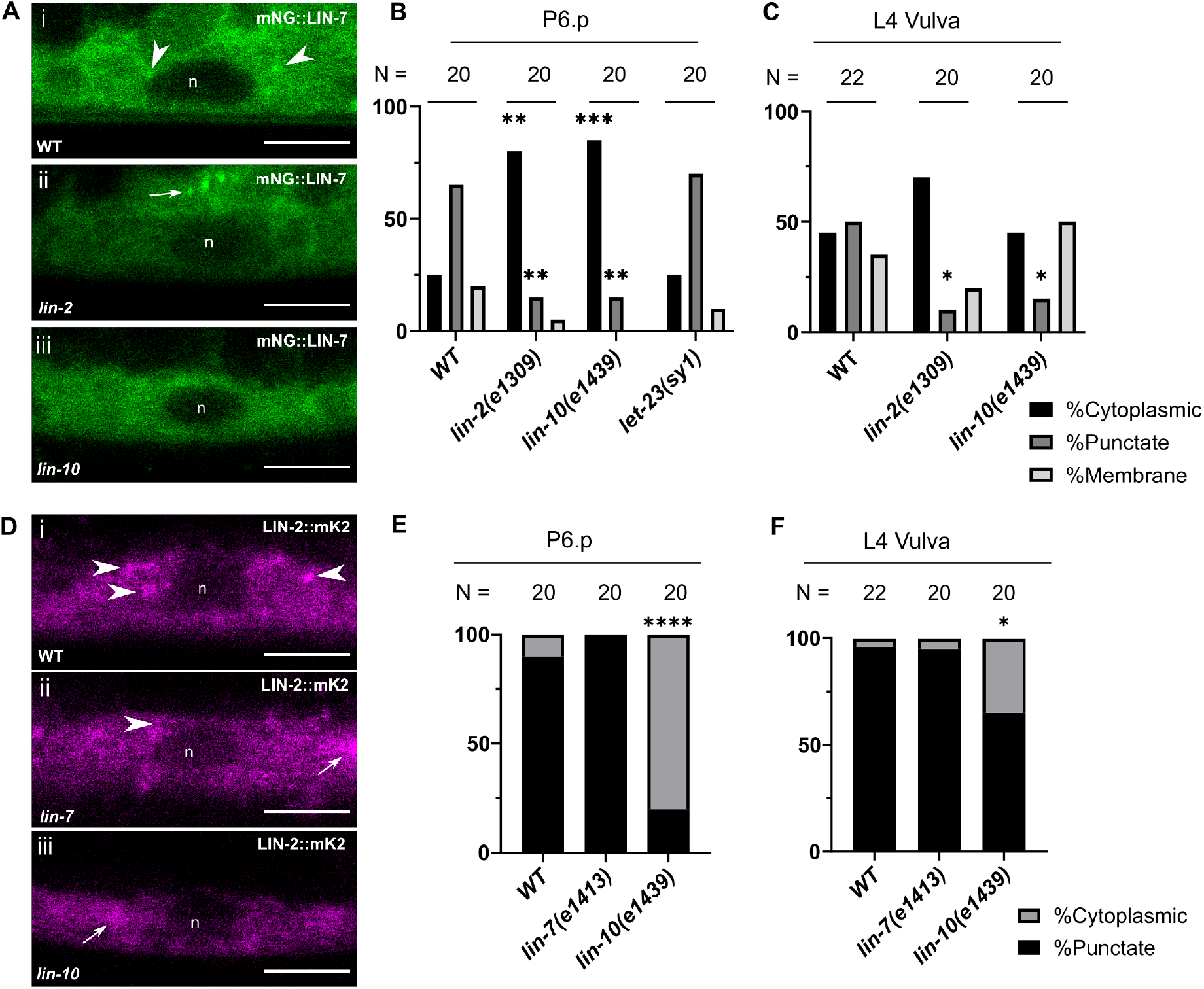
Punctate localization of LIN-2 and LIN-7 are complex-dependent. **(a-c)** Analysis of endogenously-tagged mNG::LIN-7 localization in P6.p (a,b) and L4 vulval lineages (c) in the indicated genotypes. Arrowhead: punctate localization of mNG::LIN-7 in P6.p. Arrow: punctate localization of mNG::LIN-7 in anchor cell. **(d-f)** Analysis of endogenously-tagged LIN-2::mK2 localization in P6.p (d,e) and L4 vulval lineages (f) in the indicated genotypes. Arrowhead: punctate localization of LIN-2::mK2 in P6.p. Arrow: fluorescent expression in VNC in same focal plane as VPCs. n: nucleus. N: numbers of animals scored. Scalebars: 5 μm. *p<0.05, **p<0.01, ***p<0.001, ****p<0.0001 Fisher’s Exact Test compared to WT.

We found that LIN-2 localization to cytoplasmic foci is in turn partly LIN-10-dependent, and LIN-7-independent. The predominately punctate LIN-2::mK2 becomes exclusively cytoplasmic in 80% worms at the L3 stage in a *lin-10(e1439)* null mutant (Figure 8d-e). The dependency was less pronounced, but still evident at the L4 stage (Figure 8f). Consistently, GFP::LIN-2a, expressed from an extrachromosomal transgene, localized to faint punctae in approximately 30% of VPCs imaged, was almost completely mislocalized into the cytosol in a *lin-10(e1439)* mutant (Figure S2). Punctate localization of LIN-2 was unaltered in a *lin-7(e1413)* mutant in the VPCs (Figure 8d-f).

On the other hand, we found that both endogenously-tagged mNG::LIN-10 and extrachromosomal GFP::LIN-10a punctate localization was not altered with loss-of-function mutations in *lin-2* and *lin-7* (Figure 9a-b), suggesting that LIN-10 maintains its localization pattern independently of complex formation.

**Figure 9:**
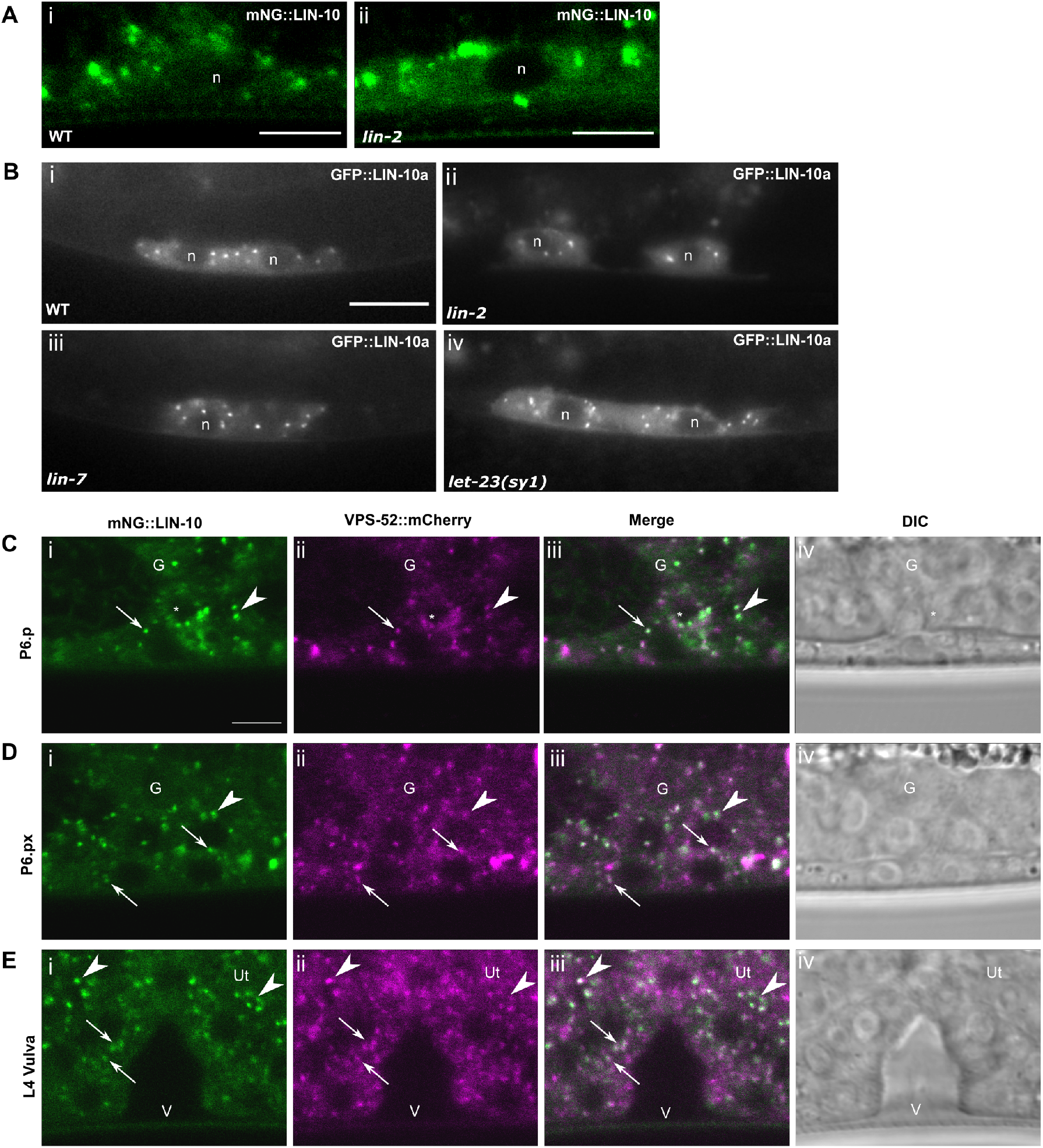
LIN-10 localization is complex-independent and overlaps with Golgi/recycling endosome protein VPS-52. **(a)** Endogenously-tagged mNG::LIN-10a localizes to cytoplasmic punctae in VPCs independent of its interacting partner LIN-2. Scalebar 5 μm. **(b)** Extrachromosomal GFP::LIN-10a localizes to punctae in the VPCs of wildtype (i) and *lin-2* (ii), *lin-7* (iii), and *let-23(sy1)* (iv) mutants. n: nucleus. Scalebar 10 μm. **(c-e)** mNG::LIN-10 colocalizes with Golgi and recycling endosome marker VPS-52 *(qbcSi01)* in VPCs (c-d) and differentiating vulval cells (e). Asterisk (*): anchor cell. G: developing gonad in L3 worms. Ut: Uterus. V: Vulval lumen. Arrow: colocalizing punctae in VPCs (a-b) and vulva (c). Arrowhead:Colocalizing punctae in gonad (a-b) and uterus (c). Scalebar: 5 μm. Images are all at same scale.

Finally, we found that both mNG::LIN-10 and mNG::LIN-7 localization was LET-23 EGFR-independent. The *sy1* allele of the *let-23 egfr* gene contains an early stop codon that truncates the last 6 amino acids, resulting in the loss of LET-23 EGFR protein interaction to the PDZ domain of LIN-7 (Aroian et al., 1994, Aroian and Sternberg, 1991, Kaech et al., 1998). Although the *sy1* mutant LET-23 is otherwise signaling-competent in other LET-23-dependent developmental events in the animal, it is localized exclusively to the apical membrane in the VPCs and cannot induce the vulval cell fate (Kaech et al., 1998). LIN-7 was not also mislocalized to apical membranes in a *let-23(sy1)* mutant; instead, LIN-7 remained associated to basolateral membranes in 20% of P6.p cells analyzed, and retained its punctate localization in 70% of cells (Figure 8b). Extrachromosomal GFP::LIN-10a localization is also unaltered in a *let-23(sy1)* mutant (Figure 9b). This suggests that mNG::LIN-7 and mNG::LIN-10 are appropriately localized to their respective subcellular domains independently of an association with LET-23 EGFR C-terminal PDZ interaction motif.

### LIN-10-positive punctae may represent Golgi mini-stacks or recycling endosomes

LIN-10 has been shown to localize to Golgi mini-stacks and recycling endosomes in *C. elegans* neurons (Glodowski et al., 2005, Whitfield et al., 1999, Zhang et al., 2012), and the homologous APBA1-3 proteins localize to the *trans-*Golgi network in mammalian neurons (Borg et al., 1999, Stricker and Huganir, 2003). To validate that punctate LIN-10 localization in *C. elegans* VPCs correspond to similar subcellular compartments, we tested for colocalization with VPS-52, a shared subunit of the *trans-*Golgi-associated GARP complex and recycling endosome-associated EARP complex (Conibear and Stevens, 2000, Liewen et al., 2005, Luo et al., 2011, Schindler et al., 2015). LIN-10-positive punctae regularly overlap with VPS-52 in the VPCs of L3 larvae and the differentiating vulval cells of L4 larvae (Figure 9c-e). LIN-10 and VPS-52 also colocalize in the surrounding tissue, such as the developing L3 gonad (including the anchor cell, the source of the LIN-3 EGF ligand) and the uterine epithelia of L4 larvae (Figure 9c-e). This observation is consistent with mNG::LIN-10 localizing to Golgi and recycling endosomes in *C. elegans* epithelial cells.

## Discussion

The evolutionarily conserved LIN-2/7/10 complex has been well characterized biochemically and genetically, but where and how it regulates LET-23 EGFR localization to the basolateral membrane of the VPCs was not clear. Here, we tagged each endogenous component of the LIN-2/7/10 complex and systematically characterized their expression and localization, their co-localization with each other and with LET-23 EGFR, and their interdependence for localization. LIN-7 was the only complex component observed to localize with LET-23 EGFR at the basolateral membrane of the VPCs. We identify intracellular compartments, likely Golgi ministacks and/or recycling endosomes, as the common site of LET-23 EGFR localization with the LIN-2/7/10 complex. Furthermore, we demonstrate that LIN-10 is largely required for recruitment of LIN-2 and LIN-7 to these intracellular compartments and point toward a role for the LIN-2/7/10 complex in basolateral membrane targeting of intracellular LET-23 EGFR.

Knowing where the LIN-2/7/10 complex forms and localizes with LET-23 EGFR provides insight into how the complex might function to promote LET-23 EGFR localization to the basolateral membrane. For example, localization to the basolateral membrane might suggest a role in anchoring LET-23 EGFR in the basolateral compartment. While LIN-7 does localize to the basolateral membrane with LET-23 EGFR, the common site of localization of the LIN-2/7/10 complex with LET-23 EGFR turns out to be intracellular compartments. LIN-10 is the most-studied of the complex components in *C. elegans* for its complex-independent role in regulating trafficking and recycling of the glutamate receptor GLR-1 in interneurons. LIN-10 has been shown to localize to Golgi and recycling endosomes in both neurons and the intestinal cells (Glodowski et al., 2005, Whitfield et al., 1999, Zhang et al., 2012). Immunolocalization of LIN-10 in the Pn.pxx cells showed a punctate localization (Whitfield et al., 1999). Mammalian LIN-10 homologs, the APBA proteins, localize to the Golgi and are involved in protein secretion, possibly functioning as a clathrin adaptor (Borg et al., 1999, Hill et al., 2003, Stricker and Huganir, 2003). Thus, it was not surprising that LIN-10 showed strong localization to intracellular compartments that overlap with VPS-52, a proposed Golgi and recycling endosome protein in the VPCs. However, we did not observe LIN-10 localization at the basolateral membrane, and LIN-10 only colocalized with LET-23 EGFR on intracellular compartments, suggesting that LIN-10 promotes basolateral secretion or recycling of LET-23 EGFR to the plasma membrane from the Golgi and/or recycling endosomes. In contrast to LIN-10, LIN-2 and LIN-7 are highly cytosolic with less pronounced localization to cytoplasmic foci that partly overlap with LIN-10 in the VPCs. Our findings that LIN-7 punctate localization is largely dependent on LIN-2 and LIN-10, that LIN-2 is dependent only on LIN-10, and that LIN-10 localization is independent of LIN-2 and LIN-7 suggests that LIN-10 recruits LIN-2, which in turn recruits LIN-7 to Golgi or recycling endosomes. These data are also consistent with previous protein interaction studies showing that LIN-2 bridges an interaction between LIN-7 and LIN-10 (Kaech et al., 1998). Coupled with our finding that LET-23 EGFR partially overlaps with LIN-10 on punctae, this suggests that the complex likely forms at the Golgi or recycling endosomes to target LET-23 EGFR to the basolateral membrane. Curiously, the overlap between LIN-10 and LIN-7 is not entirely dependent on LIN-2, indicating that the small population of LIN-7 that localizes to punctae independently of LIN-2 may occupy similar endomembrane compartments as LIN-10.

Localization to intracellular punctae is a novel finding for LIN-7 and its orthologues, and has been infrequently reported for LIN-2 and mammalian CASK (Borg et al., 1999, Tong et al., 2015). Incidentally, localization to endomembrane compartments is consistent with reported function for these two proteins. Mammalian Lin7 has been found to promote the targeting of endocytosed cargo back to the basolateral membrane (Shelly et al., 2003, Straight et al., 2001), and LIN-2 was found to interact with EPS-8, which itself localizes to cytoplasmic foci in *C. elegans* VPCs, to inhibit internalization of LET-23 EGFR to the basolateral membrane (Stetak et al., 2006). However, localization to endocytic membranes has not previously been demonstrated for LIN-2 or LIN-7. The stronger colocalization of LIN-2 and LIN-7 on intracellular foci than of either with LIN-10 suggests that LIN-2 and LIN-7 may reside on distinct compartments from LIN-10. This could suggest a LIN-10-independent function or possibly a transport intermediate.

Additionally, we found that LIN-7 uniquely localizes to the basolateral membrane, where it can partly colocalize with LET-23 EGFR, consistent with reported localization for mammalian Lin7 (Straight et al., 2000). Although we did not detect any enrichment of LIN-2 or LIN-10 at the plasma membrane, LIN-10-positive punctae can be found overlapping with LET-23 EGFR at the cell periphery. This observation may indicate that LIN-7 alone interacts with the receptor at the plasma membrane, in addition to forming a complex with LIN-2 and LIN-10 to regulate recycling or secretion of LET-23 EGFR. However, the PDZ-interaction motif of LET-23 EGFR is not required for LIN-7 localization to either punctae or the basolateral membrane, suggesting that LIN-7 localization is independent of LET-23 EGFR. Alternatively, there could be a second point of interaction between LIN-7 and LET-23 EGFR, as has been found with their mammalian orthologs (Shelly et al., 2003), that may regulate LIN-7 localization.

LIN-2 and LIN-7 are dynamically expressed during vulval development. We found that their expression levels increase in induced vulval cells during development while also decreasing in the VPCs that do not adopt vulval fates. On the other hand, LIN-10 expression remains stable throughout vulval development and in uninduced VPCs. LET-23 EGFR expression has previously been found to be upregulated in primary cell lineages (P6.p) and downregulated in secondary cell lineages (P5.p and P7.p), and MPK-1 ERK activation is amplified in primary lineages (de la Cova et al., 2017, Kaech et al., 1998, Stetak et al., 2006); however, LIN-2 and LIN-7 expression persists in both cell types. Therefore, LIN-2 and LIN-7 expression is unlikely to be regulated by LET-23 EGFR signaling. The apparent loss of LIN-2 and LIN-7 expression in uninduced VPCs might be due to diffusion of LIN-2 and LIN-7 fusion proteins across the syncytial tissue of the hypodermis following fusion of uninduced VPCs with Hyp7. The upregulation of LIN-2 and LIN-7 expression during vulval morphogenesis suggests these proteins may have other, uncharacterized roles in vulval development as suggested by previously reported non-Vul egg-laying defective phenotypes seen in hypomorphic *lin-2* and *lin-7* mutants (Ferguson and Horvitz, 1985).

Beyond the VPCs, the LIN-2/7/10 complex was found to regulate insulin signaling pathway activation and infection sensitivity in *C. elegans* through an interaction with the insulin receptor DAF-2 (Sem et al., 2012). Otherwise, the complex has not been identified in other cellular processes. LIN-2 and LIN-10 have been independently implicated in neuronal cell traffic (Rongo et al., 1998, Wu et al., 2016) and in muscle, LIN-2 localizes to neuromuscular junctions along the nerve cord (Tong et al., 2015, Zhou et al., 2020). Our findings of prominent expression of LIN-2/7/10 proteins in neuronal tissue, strong colocalization of LIN-2 and LIN-7 in the neural ring and nerve chords, and the role of the mammalian CASK/Lin7/APBA1 complex in regulating synaptic plasticity and function, suggests the complex may also form in *C. elegans* neurons. In general, LIN-2 and LIN-7 strongly colocalize with each other across most tissues and likely reflects why LIN-2 associates more strongly with LIN-7 than with LIN-10 in *in vivo* pull-down experiments. Together, these data suggest that LIN-2 interacts more transiently or infrequently with LIN-10 than with LIN-7, or that LIN-2 and LIN-7 could form a complex without LIN-10. Although not a focus of our analysis, there were two apparent tissues in which LIN-2 and LIN-7 expression differed. LIN-7 and LIN-10, but not LIN-2, were also identified in cells of the developing gonad (including the LIN-3 EGF-releasing anchor cell) and uterine epithelial cells in L3 and L4 larvae, respectively. Thus, LIN-7 and LIN-10 might have LIN-2-independent functions in the developing gonad. Conversely, LIN-2 is more strongly expressed in the isthmus of the pharynx than LIN-7.

In summary, we present the first comprehensive analysis of LIN-2, LIN-7 and LIN-10 localization in the VPCs during LET-23 EGFR-mediated vulva induction and the subsequent development of the vulva through the L4 stage. Our results are consistent with a model in which LIN-10 recruits LIN-2 and LIN-7 to Golgi ministacks or recycling endosomes to promote basolateral targeting of LET-23 EGFR. Our results also suggest that LIN-7 could have additional roles at the plasma membrane, and the upregulation of LIN-2 and LIN-7 after vulval cell fate induction hint at additional roles for these proteins at later stages of development. This study offers new insight into the localization and regulation of this evolutionarily conserved complex of scaffolding proteins, and provides a foundation to further explore their role in cell signaling regulation and development.

## Acknowledgements

We thank Tam Duong and David Reiner (Texas A&M University) for sharing unpublished strains and reagents, Martin Simard (Université Laval) and Chris Rongo (Rutgers University) for strains and reagents, Sarah Gagnon and David Reiner for comments on the manuscript, Min Fu and Shibo Feng at the RI-MUHC Molecular Imaging Platform for technical assistance, WormAtlas, and WormBase for strain information. Some strains were provided by the CGC, which is funded by NIH Office of Research Infrastructure Programs (P40 OD010440).

## Supplementary Figures and Tables

**Figure S1:**
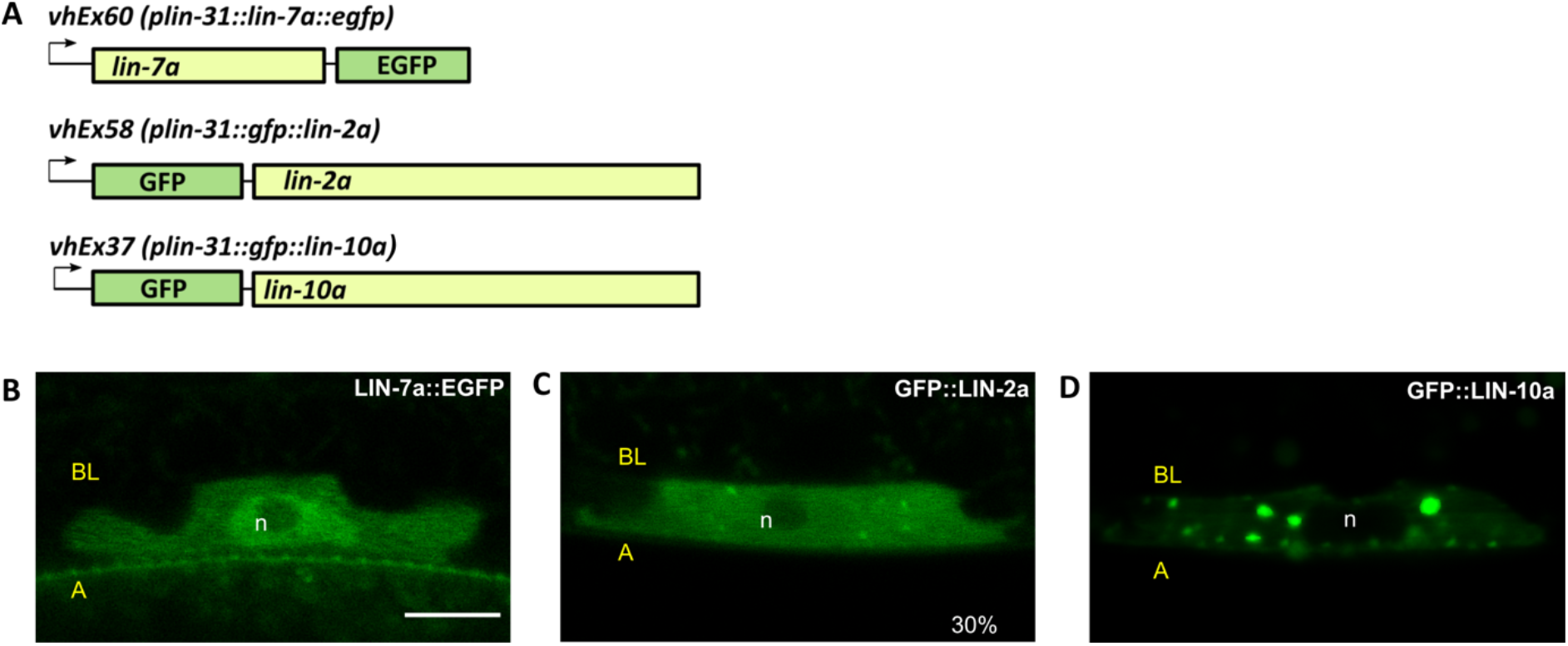
Localization of LIN-2/7/10 transgenes expressed from extrachromosomal arrays. **(a)** Schematic of extrachromosomal array transgenes *vhEx60 (plin-31::lin-7a::egfp), vhEx58 (plin-31::gfp::lin-2a),* and *vhEx37 (plin-31::gfp::lin-10a).* **(b)** LIN-7a::EGFP is exclusively cytosolic and nuclear in VPCs. Scalebar, 5 μm. **(c)** GFP::LIN-2a localizes diffusely to the cytosol and nucleus in the VPCs. In 30% of VPCs, LIN-2a also localizes to faint cytosolic foci. Scale as in (b). **(d)** GFP::LIN-10a localizes to cytoplasmic punctae and is expressed diffusely in the cytosol of VPCs. Scale as in (b). A: Apical. BL: Basolateral. n: Nucleus.

**Figure S2:**
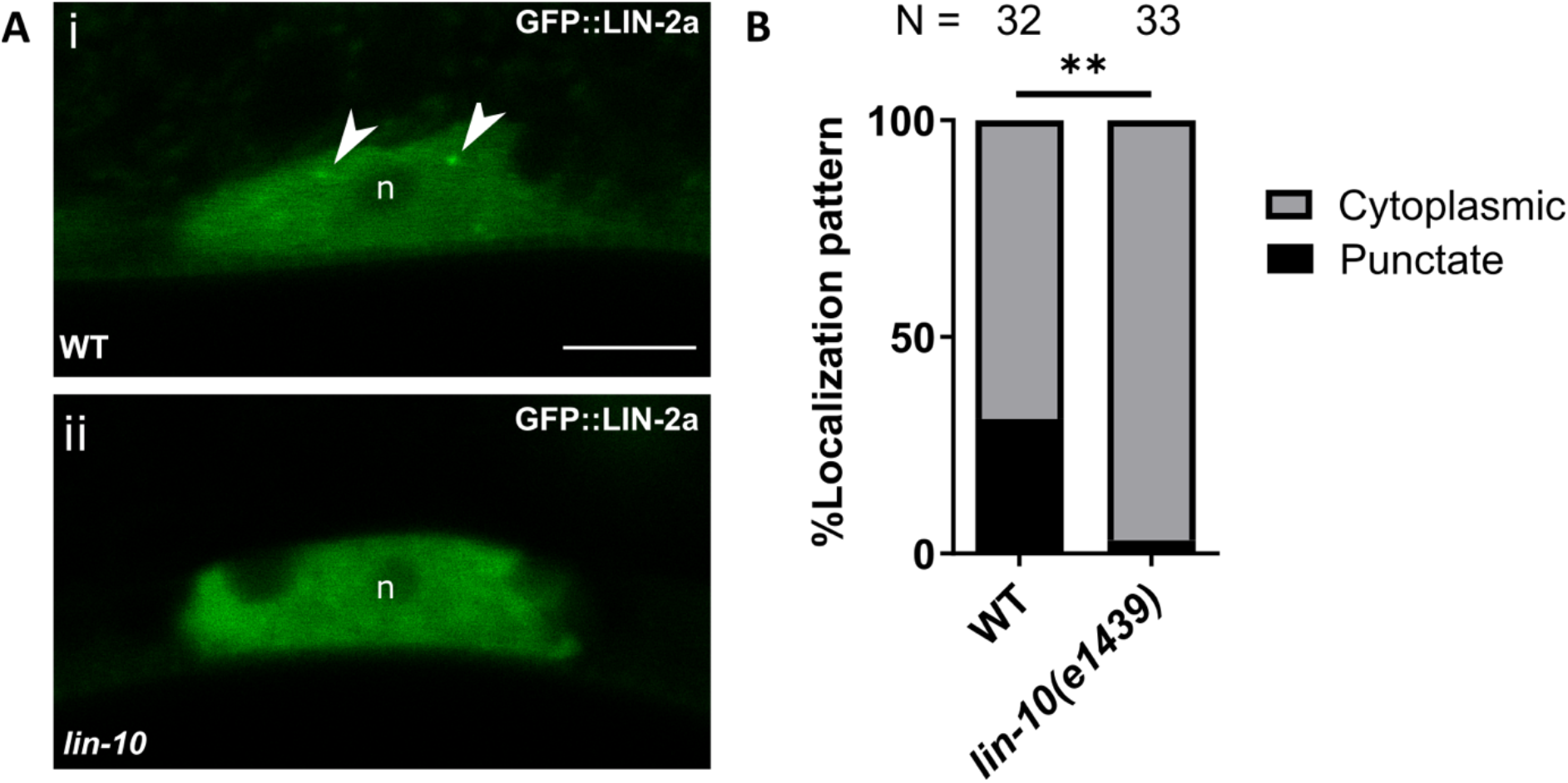
LIN-2 punctate localization is LIN-10-dependent. **(a-b)** Extrachromosomal GFP::LIN-2a localization in wildtype (a.i) and a *lin-10* mutant (a.ii), quantified in (b). Arrowheads: GFP::LIN-2a-positive punctae. n: nucleus. N, number of animals scored. Scalebars: 5 μm. Fisher’s Exact Test. **p<0.01.

**Table S1:** Strain list

| Strain name | Genotype | Source |
| --- | --- | --- |
| QR887 | *let-23(re202) lin-7(vh51) II* | This study |
| DV3366 | *let-23(re202[let-23::mKate2::3xFlag]) II* | David Reiner |
| QR930 | *let-23(sy1) lin-7(vh51) II* | This study |
| QR751 | *let-23(sy1) unc-4(e120) II; vhEx37[plin-31::gfp::lin-10a + pttx-3::gfp]* | This study |
| CB1439 | *lin-10(e1439) I* | CGC |
| QR878 | *lin-10(e1439) I; lin-2(vh52) X* | This study |
| QR929 | *lin-10(e1439) I; lin-7(vh51) II* | This study |
| QR747 | *lin-10(e1439) I; vhEx37[plin-31::gfp::lin-10a + pttx-3::gfp]* | This study |
| QR886 | *lin-10(vh50) I; let-23(re202) II* | This study |
| QR826 | *lin-10(vh50) I; lin-2(e1309) X* | This study |
| QR876 | *lin-10(vh50) I; lin-2(vh52) X* | This study |
| QR845 | *lin-10(vh50) I; qbcSi01[pvps-52::vps-52::mCherry] II* | This study |
| QR769 | *lin-10(vh50[mNeonGreen::3xFlag::LIN-10]) I* | This study |
| QR723 | *lin-2(e1309) X; vhEx37[plin-31::gfp::lin-10a + pttx-3::gfp]* | This study |
| QR724 | *lin-2(e1309) X; vhEx58[plin-31::gfp::lin-2a + pttx-3::gfp]* | This study |
| CB1309 | *lin-2(e1309) X* | CGC |
| QR830 | *lin-2(vh52[lin-2::mKate2::3xMyc]) X* | This study |
| CB1413 | *lin-7(e1413) II* | CGC |
| QR874 | *lin-7(e1413) II; lin-2(vh52) X* | This study |
| QR734 | *lin-7(e1413) II; vhEx37[plin-31::gfp::lin-10a + pttx-3::gfp]* | This study |
| QR725 | *lin-7(e1413) II; vhEx60[plin-31::lin-7a::egfp + pttx-3::gfp]* | This study |
| QR877 | *lin-7(vh51) II; lin-2(e1309) X* | This study |
| QR935 | *lin-7(vh51) II; lin-2(e1309) X; vhEx63[plin31::mCherry::lin-10a + pttx-3::gfp]* | This study |
| QR846 | *lin-7(vh51) II; lin-2(vh52) X* | This study |
| QR934 | *lin-7(vh51) II; vhEx63[plin-31::mCherry::lin-10a + pttx-3::gfp]* | This study |
| QR829 | *lin-7(vh51[mNeonGreen::3xFlag::LIN-7]) II* | This study |
| QR600 | *vhEx37[plin-31::gfp::lin-10a + pttx-3::gfp]* | This study |
| QR710 | *vhEx58[plin-31::gfp::lin-2a + pttx-3::gfp]* | This study |
| QR712 | *vhEx60[plin-31::lin-7a::egfp + pttx-3::gfp]* | This study |
| QR715 | *vhEx63[plin-31::mCherry::lin-10a + pttx-3::gfp]* | This study |
|  | *zhIs035[plet-23::let-23::gfp + unc-119(+)] I* | (Haag et al., 2014) |
| QR480 | *zhIs035[plet-23::let-23::gfp + unc-119(+)] I; lin-2(e1309) X* | (Skorobogata et al., 2014) |
| QR732 | *zhIs035[plet-23::let-23::gfp + unc-119(+)] I; lin-2(e1309) X; vhEx63[plin-31::mCherry::lin-10a + pttx-3::gfp]* | This study |
| QR727 | *zhIs035[plet-23::let-23::gfp + unc-119(+)] I; vhEx63[plin-31::mCherry::lin-10a + pttx-3::gfp]* | This study |

**Table S2:**
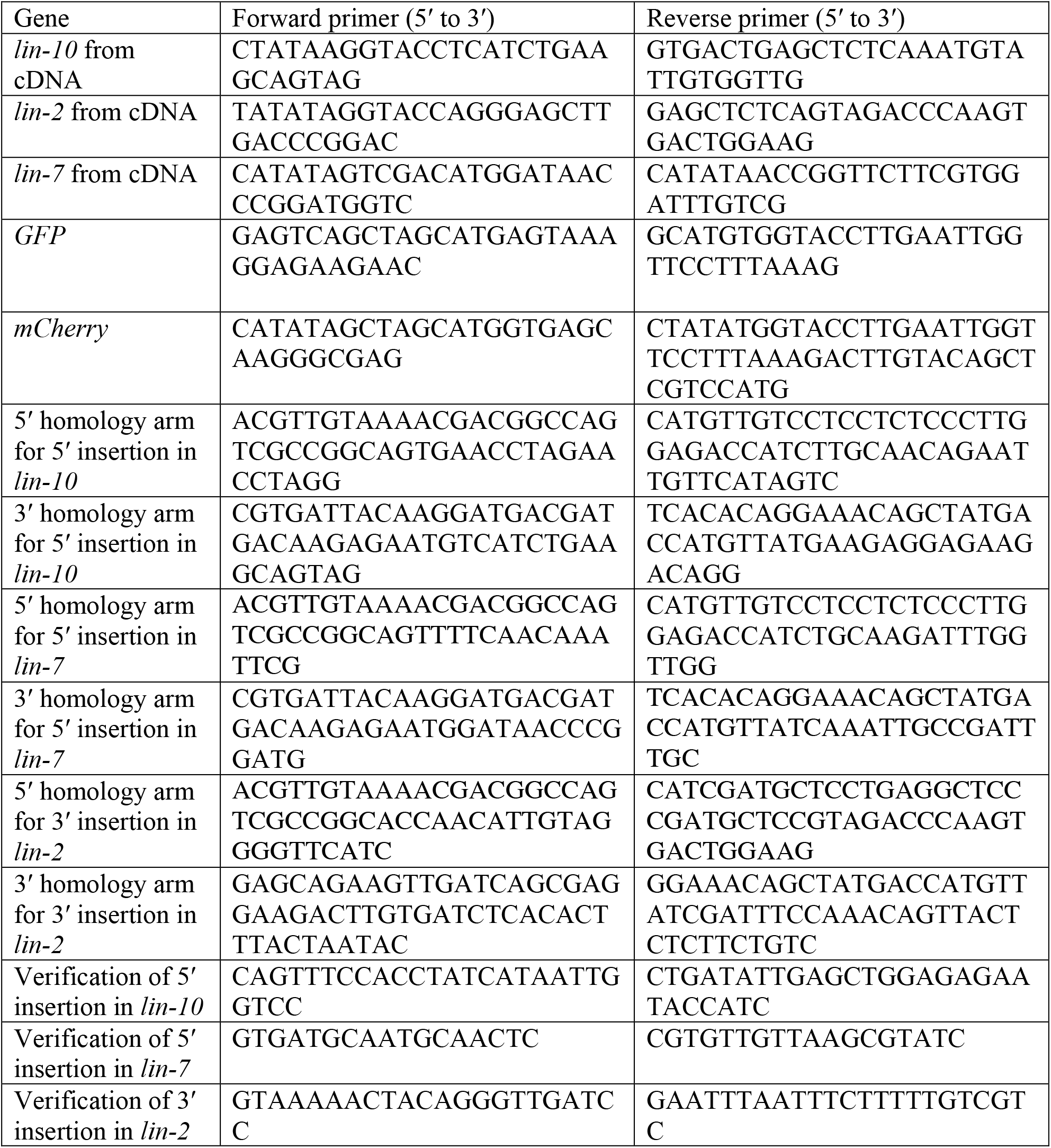
List of primers used for cloning

**Table S3:** Analysis of VPC induction in *lin-7(vh51), lin-2(vh52), lin-10(vh50),* and *let-23(re202)*

| Genotype | %Vul | %Muv | Avg # of VPCs induced | n |
| --- | --- | --- | --- | --- |
| N2 (wildtype) | 0% | 0% | 3.00 | 36 |
| *lin-7(vh51[mNG::3xFlag::LIN-7])* II | 0% | 0% | 3.00 | 47 |
| *lin-2(vh52[LIN-2::mK2::3xMyc])* X | 0% | 0% | 3.00 | 47 |
| *lin-10(vh50[mNG::3xFlag::LIN-10])* I | 0% | 0% | 3.00 | 45 |
| *let-23(re202[LET-23::mK2::3xFlag])* II | 5% | 7% | 3.02 | 42 |

One-way ANOVA for VPC induction with Dunnett's test for multiple comparisons. Fisher's exact test for Vul and Muv phenotypes. All conditions compared to N2 (wildtype).

**Table S4:** Extrachromosomal LIN-7a::EGFP, GFP::LIN-2a, and GFP::LIN-10a rescue their respective mutant phenotypes

| Genotype | %Vul | %Muv | Avg # of VPCs induced | n |
| --- | --- | --- | --- | --- |
| 1 *lin-7(e1413)* | 82% | 0% | 1.15 | 17 |
| 2 *lin-7(e1413); vhEx60(plin-31::lin-7a::egfp)* | 27%** | 7% | 2.83**** | 15 |
| 3 *lin-2(e1309)* | 96% | 0% | 0.39 | 23 |
| 4 *lin-2(e1309); vhEx58(plin-31::gfp::lin-2a)* | 8%**** | 8% | 2.92**** | 13 |
| 5 *lin-10(e1439)* | 91% | 0% | 0.87 | 22 |
| 6 *lin-10(e1439); vhEx37(plin-31::gfp::lin-10a)* | 8%**** | 6% | 2.94**** | 36 |

Two-tailed Student's *t*-test. Fisher's exact test for Vul and Muv phenotypes compared to the shaded row. \*\*P<0.01, \*\*\*\*P<0.0001. Row 2 was compared with row 1. Row 4 was compared with row 3. Row 6 was compared with row 5.

## References

Abdus-Saboor, I., Mancuso, V. P., Murray, J. I., Palozola, K., Norris, C., Hall, D. H., Howell, K., Huang, K. & Sundaram, M. V. (2011). Notch and Ras promote sequential steps of excretory tube development in C. *elegans*. Development, 138, 3545–55.

Alewine, C., Kim, B. Y., Hegde, V. & Welling, P. A. (2007). Lin-7 targets the Kir 2.3 channel on the basolateral membrane via a L27 domain interaction with CASK. Am J Physiol Cell Physiol, 293, C1733–41.

Aroian, R. V., Lesa, G. M. & Sternberg, P. W. (1994). Mutations in the Caenorhabditis elegans let-23 EGFR-like gene define elements important for cell-type specificity and function. EMBO J, 13, 360–6.

Aroian, R. V. & Sternberg, P. W. (1991). Multiple functions of let-23, a Caenorhabditis elegans receptor tyrosine kinase gene required for vulval induction. Genetics, 128, 251–67.

Becamel, C., Alonso, G., Galeotti, N., Demey, E., Jouin, P., Ullmer, C., Dumuis, A., Bockaert, J. & Marin, P. (2002). Synaptic multiprotein complexes associated with 5-HT(2C) receptors: a proteomic approach. EMBO J, 21, 2332–42.

Borg, J. P., Lopez-Figueroa, M. O., de Taddeo-Borg, M., Kroon, D. E., Turner, R. S., Watson, S. J. & Margolis, B. (1999). Molecular analysis of the X11-mLin-2/CASK complex in brain. J Neurosci, 19, 1307–16.

Borg, J. P., Straight, S. W., Kaech, S. M., de Taddeo-Borg, M., Kroon, D. E., Karnak, D., Turner, R. S., Kim, S. K. & Margolis, B. (1998). Identification of an Evolutionarily Conserved Heterotrimeric Protein Complex Involved in Protein Targeting. Journal of Biological Chemistry, 273, 31633–31636.

Brenner, S. (1974). The genetics of Caenorhabditis elegans. Genetics, 77, 71–94.

Buday, L. & Tompa, P. (2010). Functional classification of scaffold proteins and related molecules. FEBS J, 277, 4348–55.

Butz, S., Okamoto, M. & Sudhof, T. C. (1998). A tripartite protein complex with the potential to couple synaptic vesicle exocytosis to cell adhesion in brain. Cell, 94, 773–82.

Chen, K. & Featherstone, D. E. (2011). Pre and postsynaptic roles for Drosophila CASK. Mol Cell Neurosci, 48, 171–82.

Cohen, A. R., Woods, D. F., Marfatia, S. M., Walther, Z., Chishti, A. H. & Anderson, J. M. (1998). Human CASK/LIN-2 binds syndecan-2 and protein 4.1 and localizes to the basolateral membrane of epithelial cells. J Cell Biol, 142, 129–38.

Conibear, E. & Stevens, T. H. (2000). Vps52p, Vps53p, and Vps54p form a novel multisubunit complex required for protein sorting at the yeast late Golgi. Mol Biol Cell, 11, 305–23.

Cristofoli, F., Devriendt, K., Davis, E. E., Van Esch, H. & Vermeesch, J. R. (2018). Novel CASK mutations in cases with syndromic microcephaly. Hum Mutat, 39, 993–1001.

de la Cova, C., Townley, R., Regot, S. & Greenwald, I. (2017). A Real-Time Biosensor for ERK Activity Reveals Signaling Dynamics during C. *elegans* Cell Fate Specification. Dev Cell, 42, 542–553 e4.

Dickinson, D. J., Pani, A. M., Heppert, J. K., Higgins, C. D. & Goldstein, B. (2015). Streamlined Genome Engineering with a Self-Excising Drug Selection Cassette. Genetics, 200, 1035–49.

Fairless, R., Masius, H., Rohlmann, A., Heupel, K., Ahmad, M., Reissner, C., Dresbach, T. & Missler, M. (2008). Polarized targeting of neurexins to synapses is regulated by their C-terminal sequences. J Neurosci, 28, 12969–81.

Ferguson, E. L. & Horvitz, H. R. (1985). Identification and characterization of 22 genes that affect the vulval cell lineages of the nematode Caenorhabditis elegans. Genetics, 110, 17–72.

Gauthier, K. & Rocheleau, C. E. (2017). C. elegans Vulva Induction: An In Vivo Model to Study Epidermal Growth Factor Receptor Signaling and Trafficking. Methods Mol Biol, 1652, 43–61.

Glodowski, D. R., Wright, T., Martinowich, K., Chang, H. C., Beach, D. & Rongo, C. (2005). Distinct LIN-10 domains are required for its neuronal function, its epithelial function, and its synaptic localization. Mol Biol Cell, 16, 1417–26.

Gruel, N., Fuhrmann, L., Lodillinsky, C., Benhamo, V., Mariani, O., Cedenot, A., Arnould, L., Macgrogan, G., Sastre-Garau, X., Chavrier, P., et al. (2016). LIN7A is a major determinant of cell-polarity defects in breast carcinomas. Breast Cancer Res, 18, 23.

Haag, A., Gutierrez, P., Buhler, A., Walser, M., Yang, Q., Langouet, M., Kradolfer, D., Frohli, E., Herrmann, C. J., Hajnal, A., et al. (2014). An in vivo EGF receptor localization screen in C. *elegans* Identifies the Ezrin homolog ERM-1 as a temporal regulator of signaling. PLoS Genet, 10, e1004341.

Hara, T., Murakami, Y., Seiki, M. & Sakamoto, T. (2017). Mint3 in bone marrow-derived cells promotes lung metastasis in breast cancer model mice. Biochem Biophys Res Commun, 490, 688–692.

Harlow, E. & Lane, D. (2006). Stripping immunoblots for reprobing or storage. CSH Protoc, 2006.

Hill, K., Li, Y., Bennett, M., McKay, M., Zhu, X., Shern, J., Torre, E., Lah, J. J., Levey, A. I. & Kahn, R. A. (2003). Munc18 interacting proteins: ADP-ribosylation factor-dependent coat proteins that regulate the traffic of beta-Alzheimer’s precursor protein. J Biol Chem, 278, 36032–40.

Hirose, Y., Johnson, Z. I., Schoepflin, Z. R., Markova, D. Z., Chiba, K., Toyama, Y., Shapiro, I. M. & Risbud, M. V. (2014). FIH-1-Mint3 axis does not control HIF-1 transcriptional activity in nucleus pulposus cells. J Biol Chem, 289, 20594–605.

Hobert, O., Mori, I., Yamashita, Y., Honda, H., Ohshima, Y., Liu, Y. & Ruvkun, G. (1997). Regulation of interneuron function in the C. *elegans* thermoregulatory pathway by the ttx-3 LIM homeobox gene. Neuron, 19, 345–57.

Hsueh, Y. P. (2006). The role of the MAGUK protein CASK in neural development and synaptic function. Curr Med Chem, 13, 1915–27.

Hsueh, Y. P., Wang, T. F., Yang, F. C. & Sheng, M. (2000). Nuclear translocation and transcription regulation by the membrane-associated guanylate kinase CASK/LIN-2. Nature, 404, 298–302.

Hung, M. C. & Link, W. (2011). Protein localization in disease and therapy. J Cell Sci, 124, 3381–92.

Jeyifous, O., Waites, C. L., Specht, C. G., Fujisawa, S., Schubert, M., Lin, E. I., Marshall, J., Aoki, C., de Silva, T., Montgomery, J. M., et al. (2009). SAP97 and CASK mediate sorting of NMDA receptors through a previously unknown secretory pathway. Nat Neurosci, 12, 1011–9.

Jo, K., Derin, R., Li, M. & Bredt, D. S. (1999). Characterization of MALS/Velis-1, −2, and - 3: a family of mammalian LIN-7 homologs enriched at brain synapses in association with the postsynaptic density-95/NMDA receptor postsynaptic complex. J Neurosci, 19, 4189–99.

Kaech, S. M., Whitfield, C. W. & Kim, S. K. (1998). The LIN-2/LIN-7/LIN-10 complex mediates basolateral membrane localization of the C. *elegans* EGF receptor LET-23 in vulval epithelial cells. Cell, 94, 761–71.

LaConte, L. E., Chavan, V. & Mukherjee, K. (2014). Identification and glycerol-induced correction of misfolding mutations in the X-linked mental retardation gene CASK. PLoS One, 9, e88276.

Lau, K. F., Perkinton, M. S., Rodriguez, L., McLoughlin, D. M. & Miller, C. C. (2010). An X11alpha/FSBP complex represses transcription of the GSK3beta gene promoter. Neuroreport, 21, 761–6.

Leonoudakis, D., Conti, L. R., Radeke, C. M., McGuire, L. M. & Vandenberg, C. A. (2004). A multiprotein trafficking complex composed of SAP97, CASK, Veli, and Mint1 is associated with inward rectifier Kir2 potassium channels. J Biol Chem, 279, 19051–63.

Liewen, H., Meinhold-Heerlein, I., Oliveira, V., Schwarzenbacher, R., Luo, G., Wadle, A., Jung, M., Pfreundschuh, M. & Stenner-Liewen, F. (2005). Characterization of the human GARP (Golgi associated retrograde protein) complex. Exp Cell Res, 306, 24–34.

Lozovatsky, L., Abayasekara, N., Piawah, S. & Walther, Z. (2009). CASK deletion in intestinal epithelia causes mislocalization of LIN7C and the DLG1/Scrib polarity complex without affecting cell polarity. Mol Biol Cell, 20, 4489–99.

Luo, L., Hannemann, M., Koenig, S., Hegermann, J., Ailion, M., Cho, M. K., Sasidharan, N., Zweckstetter, M., Rensing, S. A. & Eimer, S. (2011). The Caenorhabditis elegans GARP complex contains the conserved Vps51 subunit and is required to maintain lysosomal morphology. Mol Biol Cell, 22, 2564–78.

Malik, B. R. & Hodge, J. J. (2014). CASK and CaMKII function in Drosophila memory. Front Neurosci, 8, 178.

Mello, C. C., Kramer, J. M., Stinchcomb, D. & Ambros, V. (1991). Efficient gene transfer in C.elegans: extrachromosomal maintenance and integration of transforming sequences. EMBO J, 10, 3959–70.

Miller, C. C., McLoughlin, D. M., Lau, K. F., Tennant, M. E. & Rogelj, B. (2006). The X11 proteins, Abeta production and Alzheimer’s disease. Trends Neurosci, 29, 280–5.

Motodate, R., Saito, Y., Hata, S. & Suzuki, T. (2016). Expression and localization of X11 family proteins in neurons. Brain Res, 1646, 227–234.

Mugabo, Y. & Lim, G. E. (2018). Scaffold Proteins: From Coordinating Signaling Pathways to Metabolic Regulation. Endocrinology, 159, 3615–3630.

Rongo, C., Whitfield, C. W., Rodal, A., Kim, S. K. & Kaplan, J. M. (1998). LIN-10 is a shared component of the polarized protein localization pathways in neurons and epithelia. Cell, 94, 751–9.

Schindler, C., Chen, Y., Pu, J., Guo, X. & Bonifacino, J. S. (2015). EARP is a multisubunit tethering complex involved in endocytic recycling. Nat Cell Biol, 17, 639–50.

Sem, X., Kreisberg, J. F., Kawli, T., Tan, M. W., Rhen, M. & Tan, P. (2012). Modulation of Caenorhabditis elegans infection sensitivity by the LIN-7 cell junction protein. Cell Microbiol, 14, 1584–99.

Setou, M., Nakagawa, T., Seog, D. H. & Hirokawa, N. (2000). Kinesin superfamily motor protein KIF17 and mLin-10 in NMDA receptor-containing vesicle transport. Science, 288, 1796–802.

Sharma-Kishore, R., White, J. G., Southgate, E. & Podbilewicz, B. (1999). Formation of the vulva in Caenorhabditis elegans: a paradigm for organogenesis. Development, 126, 691–9.

Shelly, M., Mosesson, Y., Citri, A., Lavi, S., Zwang, Y., Melamed-Book, N., Aroeti, B. & Yarden, Y. (2003). Polar expression of ErbB-2/HER2 in epithelia. *Bimodal regulation by Lin-7*. Dev Cell, 5, 475–86.

Simske, J. S., Kaech, S. M., Harp, S. A. & Kim, S. K. (1996). LET-23 receptor localization by the cell junction protein LIN-7 during C. *elegans* vulval induction. Cell, 85, 195204.

Skorobogata, O., Escobar-Restrepo, J. M. & Rocheleau, C. E. (2014). An AGEF-1/Arf GTPase/AP-1 ensemble antagonizes LET-23 EGFR basolateral localization and signaling during C. *elegans* vulva induction. PLoS Genet, 10, e1004728.

Skorobogata, O., Meng, J., Gauthier, K. & Rocheleau, C. E. (2016). Dynein-mediated trafficking negatively regulates LET-23 EGFR signaling. Mol Biol Cell.

Sternberg, P. W. & Horvitz, H. R. (1986). Pattern formation during vulval development in C. *elegans*. Cell, 44, 761–72.

Stetak, A., Hoier, E. F., Croce, A., Cassata, G., Di Fiore, P. P. & Hajnal, A. (2006). Cell fate-specific regulation of EGF receptor trafficking during Caenorhabditis elegans vulval development. EMBO J, 25, 2347–57.

Straight, S. W., Chen, L., Karnak, D. & Margolis, B. (2001). Interaction with mLin-7 alters the targeting of endocytosed transmembrane proteins in mammalian epithelial cells. Mol Biol Cell, 12, 1329–40.

Straight, S. W., Karnak, D., Borg, J. P., Kamberov, E., Dare, H., Margolis, B. & Wade, J. B. (2000). mLin-7 is localized to the basolateral surface of renal epithelia via its NH(2) terminus. Am J Physiol Renal Physiol, 278, F464–75.

Stricker, N. L. & Huganir, R. L. (2003). The PDZ domains of mLin-10 regulate its transGolgi network targeting and the surface expression of AMPA receptors. Neuropharmacology, 45, 837–848.

Sulston, J. E. & Horvitz, H. R. (1977). Post-embryonic cell lineages of the nematode, Caenorhabditis elegans. Dev Biol, 56, 110–56.

Sumioka, A., Saito, Y., Sakuma, M., Araki, Y., Yamamoto, T. & Suzuki, T. (2008). The X11L/X11beta/MINT2 and X11L2/X11gamma/MINT3 scaffold proteins shuttle between the nucleus and cytoplasm. Exp Cell Res, 314, 1155–62.

Swistowski, A., Zhang, Q., Orcholski, M. E., Crippen, D., Vitelli, C., Kurakin, A. & Bredesen, D. E. (2009). Novel mediators of amyloid precursor protein signaling. J Neurosci, 29, 15703–12.

Tong, X. J., Hu, Z., Liu, Y., Anderson, D. & Kaplan, J. M. (2015). A network of autism linked genes stabilizes two pools of synaptic GABA(A) receptors. Elife, 4, e09648.

Wei, J. L., Fu, Z. X., Fang, M., Zhou, Q. Y., Zhao, Q. N., Guo, J. B., Lu, W. D. & Wang, H. (2014). High expression of CASK correlates with progression and poor prognosis of colorectal cancer. Tumour Biol, 35, 9185–94.

Whitfield, C. W., Benard, C., Barnes, T., Hekimi, S. & Kim, S. K. (1999). Basolateral localization of the Caenorhabditis elegans epidermal growth factor receptor in epithelial cells by the PDZ protein LIN-10. Mol Biol Cell, 10, 2087–100.

Wu, G. H., Muthaiyan Shanmugam, M., Bhan, P., Huang, Y. H. & Wagner, O. I. (2016). Identification and Characterization of LIN-2(CASK) as a Regulator of Kinesin-3 UNC-104(KIF1A) Motility and Clustering in Neurons. Traffic, 17, 891–907.

Yochem, J., Sundaram, M. & Han, M. (1997). Ras is required for a limited number of cell fates and not for general proliferation in Caenorhabditis elegans. Mol Cell Biol, 17, 2716–22.

Zhang, D., Isack, N. R., Glodowski, D. R., Liu, J., Chen, C. C., Xu, X. Z., Grant, B. D. & Rongo, C. (2012). RAB-6.2 and the retromer regulate glutamate receptor recycling through a retrograde pathway. J Cell Biol, 196, 85–101.

Zhou, X., Gueydan, M., Jospin, M., Ji, T., Valfort, A., Pinan-Lucarre, B. & Bessereau, J. L. (2020). The netrin receptor UNC-40/DCC assembles a postsynaptic scaffold and sets the synaptic content of GABAA receptors. Nat Commun, 11, 2674.

Zucker, B., Kama, J. A., Kuhn, A., Thu, D., Orlando, L. R., Dunah, A. W., Gokce, O., Taylor, D. M., Lambeck, J., Friedrich, B., et al. (2010). Decreased Lin7b expression in layer 5 pyramidal neurons may contribute to impaired corticostriatal connectivity in huntington disease. J Neuropathol Exp Neurol, 69, 880–95.

## References

Haag, A., Gutierrez, P., Buhler, A., Walser, M., Yang, Q., Langouet, M., Kradolfer, D., Frohli, E., Herrmann, C. J., Hajnal, A., et al. (2014). An in vivo EGF receptor localization screen in C. elegans Identifies the Ezrin homolog ERM-1 as a temporal regulator of signaling. PLoS Genet, 10, e1004341.

Skorobogata, O., Escobar-Restrepo, J. M. & Rocheleau, C. E. (2014). An AGEF-1/Arf GTPase/AP-1 ensemble antagonizes LET-23 EGFR basolateral localization and signaling during C. elegans vulva induction. PLoS Genet, 10, e1004728.

